# Unravelling cotton RNAseq repositories to the fiber development specific modules and their alliance with the fiber-related traits

**DOI:** 10.1101/2021.02.13.431059

**Authors:** Priti Prasad, Uzma Khatoon, Rishi Kumar Verma, Ajay Kumar, Debashish Mohapatra, Parthasarthi Bhattacharya, Sumit K Bag, Samir V Sawant

## Abstract

Cotton fiber development is still an intriguing question to understand the fiber commitment and development. Here, we remapped >350 publicly available cotton RNA sequencing data on recently published cotton genome with ∼400 fold coverage. The differentially expressed genes were clustered in six modules whose functions are specific to commitment, initiation, elongation and Secondary Cell Wall (SCW) fiber development stages. Gene Ontology analysis of commitment and initiation specific modules suggests enrichment of genes involved in organ development. The modules specific for elongation and SCW showed significant enrichment of hydroxyproline-rich proteins and hydrolases. Transcription factors (TFs) binding frequency of defined modules suggested that homeodomain, MYB and NAC expresses at commitment stages but their expression was governed by other TFs. We also mined the stage-specific transcriptional biomarker and Exclusively Expressed Transcripts (EETs) for fiber. These EETs were positively selected during fiber evolution and cotton domestication. The extensive expression profiling of six EETs in 100 cotton genotypes at different fiber developmental stages using nCounter assay and their correlation with eight fiber-related suggests that several EETs are correlated with different fiber quality-related traits. Thus, our study reveals several important genes and pathways that may be important for cotton fiber development and future improvement of cotton.

## Introduction

Transcriptome is the repertoire for protein-coding and non-coding RNAs of different tissues which are crucial in delineating the molecular basis of any phenotypic plasticity. Due to low-cost Next Generation Sequencing technologies and unprecedented sensitivity, RNA-sequencing (RNAseq) techniques are used routinely to scrutinize the genome-wide transcriptional activity. It provides precise information about the expression level of transcripts and their isoform at a single base-pair resolution (Wang et al., 2009). Since the last decade, more than 175 thousand RNAseq data of plants have been deposited in Sequence Read Archive (SRA) alone. To make sense of such public archives, reusability and reproducibility of data were highly recommended for providing new biological insight (Rung and Brazma, 2013). Many studies were conducted in this direction like genome upgradation (Cheng et al., 2017) and network interpretation in Arabidopsis (He et al., 2016), potential resistance gene identification in tomato (Torres-Avilés et al., 2014) and different integrative approaches in different plants (Mercatelli et al., 2020; Napolitano et al., 2020; Xie et al., 2020).

Cotton is one of the most important natural fiber crops which is cultivated in more than 70 countries around the world (Chen et al., 2007). Biologically, fibers are single-cell seed trichome that originates on the ovule surface and thus provides a model system for studying commitment, cell elongation and cell wall deposition (Kim and Triplett, 2001). Fiber development is a complex process that initiates at the outer integument of the ovule before the day of anthesis. It involves four distinguished and overlapping stages viz., initiation, elongation, secondary cell wall biosynthesis and maturation. The genus *Gossypium* is composed of 45 diploid and five tetraploid species. Tetraploid cotton species are natural allotetraploid formed by interspecific hybridization of A and D diploid genome followed by polyploidization. The five polyploid cotton genomes have conserved gene content with the same chromosomal orientation relative to the respective diploid genome (Chen et al., 2020). Among the polyploid cotton, *Gossypium hirsutum* is the most cultivated species, contributed 95% of total global cotton production signifies its natural adaptability. The significant evolutionary pressure involving genetic and epigenetic modifications leading to dynamic changes in the cotton genome (Paterson et al., 2012; Guan et al., 2014; Song et al., 2017; Li et al., 2019) which gradually responsible for the domestications of modern-day cotton.

Cotton fiber yield and their quality are essential agronomical traits, and thereby significant efforts have been made to understand the biological mechanism for cotton fiber development. The two independent groups sequenced the *Gossypium hirsutum* genome in 2015 and provided a draft reference (Li et al., 2015; Zhang et al., 2015) that revolutionized cotton research and a major leap in the direction of understanding cotton biology. In recent years, resequencing efforts of *Gossypium hirsutum* (Hu et al., 2019; Wang et al., 2019; Yang et al., 2019; Chen et al., 2020; Huang et al., 2020) have also significantly improvised the reference genome. However, before the cotton genome sequencing projects, significant efforts have been seen using EST sequencing (Arpat et al., 2004; Wilkins and Arpat, 2005; Samuel Yang et al., 2006; Udall et al., 2006; Taliercio and Boykin, 2007) and microarray analysis (Shi et al., 2006; Udall et al., 2007; Liu et al., 2012; Gilbert et al., 2013; Salih et al., 2016) to unravel the molecular mechanisms of cotton fiber development. The RNAseq efforts to understand the fiber development in cotton can be broadly categorized into before the availability of cotton genome (Wang et al., 2010; Nigam et al., 2014; Peng et al., 2014; Cheng et al., 2016; Parekh et al., 2018) and post-sequencing (Zhang et al., 2015; Ma et al., 2016; Fang et al., 2017; Ma et al., 2018; Qin et al., 2019). However, no coordinated efforts took into account where publically available RNAseq data and the power of fine reference genome map were used to mine the conserved features that might be important for cotton fiber development.

In our present work, with the availability of the high-quality reference genome of *Gossypium hirsutum* (Wang et al., 2019), we strengthen our understanding at the transcript level by reusing the publicly available cotton transcriptome data. We processed the 356 RNAseq data and clustered them on different fiber developmental stage modules. Our comprehensive computational effort leads to identifying stage-specific transcriptional biomarkers and Exclusively Expressed Transcripts (EETs) in various fiber development stages. Many of the identified genes were validated by correlating their expression to different fiber quality traits in 100 genotypes, and thus substantiating their importance in fiber development.

## Results

### A comprehensive gene expression atlas of the fiber development stages of cotton

A total of 40 SRA accessions (**Data S1)** were accessed and processed into 356 fastq files through the SRA toolkit (**Figure 1a**). The RNAseq dataset comprised of different cotton tissues such as anther, boll, bud, calycle, corolla, cotyledon, fiber, flower, leaf, meristem, ovule, pericarp, pistil, root, seed, stamen, stem and torus of the different cultivars (**Figure S1**). We considered data only from five tissues (Ovule, Fiber, Leaf, Root, and Seed) those having the highest number of SRA samples (**Figure 1b**). Out of approximately eight billion total reads, 6.1 billion reads were uniquely mapped to the cotton genome (**Figure 1c**) representing 400X coverage to ∼2.8 Gb genome size. We also independently processed the ∼0.8 million unmapped reads and assembled them into a super-transcripts on which 40% unmapped reads were realigned (**Figure S2**). Total 88 super-transcripts have Reads Per Kilobase Million (RPKM) value >1 and Gene Ontology (GO) and Kyoto Encyclopedia of Genes and Genomes (KEGG) pathway analysis indicates their role in metabolic pathways and oxidative phosphorylation, as well as, associated with nutrient reservoir and seed maturation activity (**Figure S3**).

**Figure 1.**
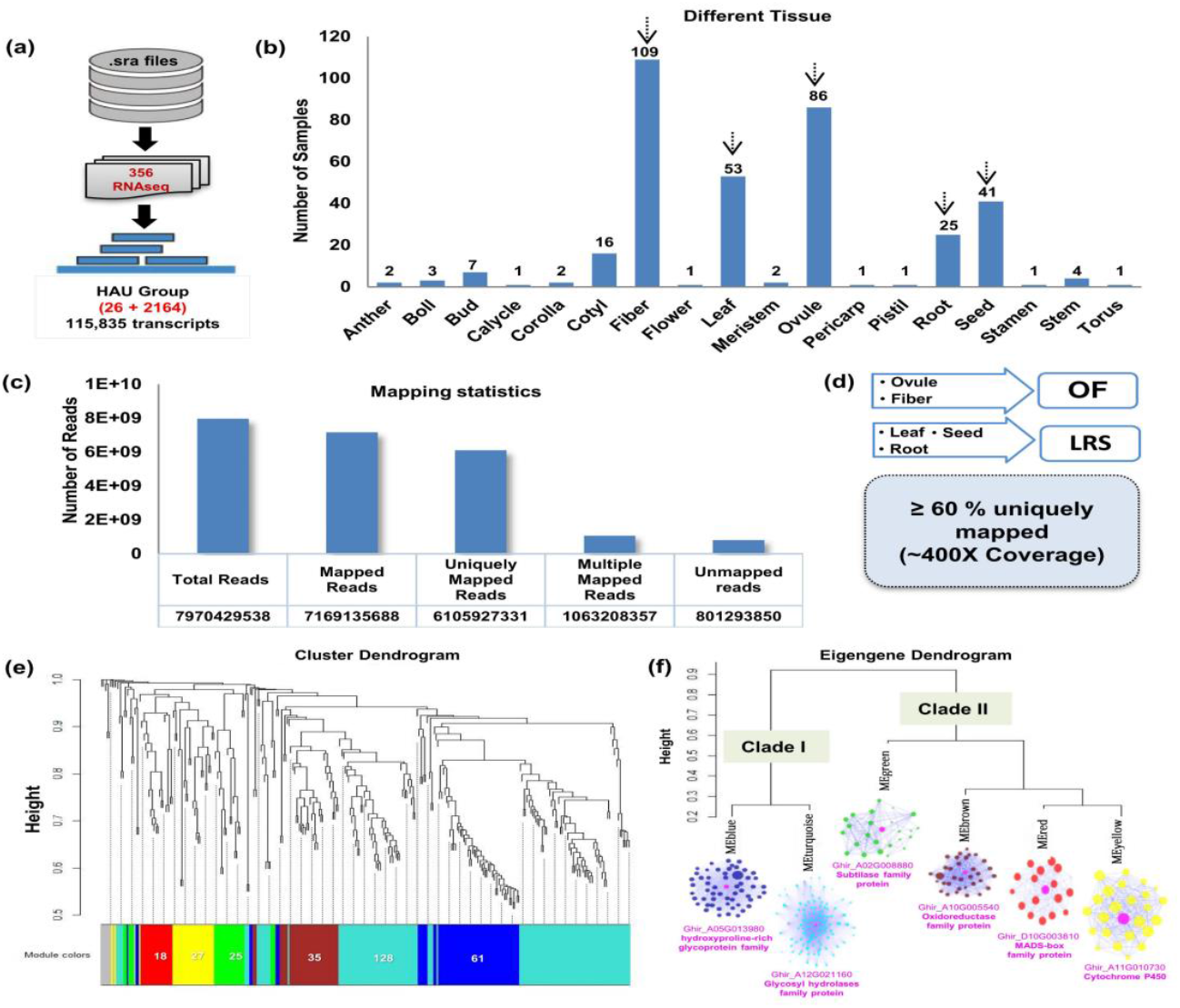
Processing of RNAseq data and their module wise clustering in OF tissues. (a) Preprocessing and mapping of the SRA files on HAU assembled genome that comprised of 115,835 transcripts. (b) Different numbers of processed RNAseq samples of different eighteen cotton tissues. The highest number of samples was indicated with the downwards arrow. (c) Mapping statistics of five selected tissues including the total number of reads, uniquely mapped reads and unmapped reads. (d) Differential estimation of uniquely mapped reads of OF and LRS tissues. OF tissue span ovule and fiber uniquely mapping reads while LRS tissue consists of uniquely mapped reads of leaf, root and seed. (e) Module-specific hierarchical clustering of up-regulated genes (302 genes with the log2fc > 2 && FDR < 0.005) in OF tissue. Different color represents different module and the encrypted number indicates the total number of genes present in that module. (f) Eigengene dendrogram of six modules that were grouped into two major clades. The co-expression behaviour of genes present in each module was visualized through a co-expression network in which magenta color shows the hub gene. The putative annotation of the hub gene was mentioned below the clade.

The primary aim of this study is to develop a comprehensive atlas of gene expression during different fiber development stages by utilizing deposited RNAseq data from extracted fiber, especially on later stages (109 samples) or fiber development on the ovules (86 samples). We pulled uniquely mapped reads (**Data S2**) from the Ovule and Fiber tissue into one group and termed it as OF, while uniquely mapped reads from Leaf, Root and Seeds were pulled together and termed it as LRS (**Figure 1d**) (**Data S3**). We identified a total of 973 differentially expressed genes (False Discovery Rate (FDR) < 0.005 and log2fold change > 2) between OF and LRS, out of which 302 genes were upregulated and 671 genes were downregulated (**Data S4**). Upregulated genes were further considered for the cluster analysis in which 8 genes were excluded as outliers (represent in grey color in **Figure 1e**). Remaining 294 genes were clustered with the soft thresholding power of 10 and constructed a blockwise module (**Figure 1e**). As a result, we obtained six modules represented in blue (61 genes), turquoise (128 genes), green (25 genes), brown (35 genes), red (18 genes) and yellow (27 genes) colored schemes as mentioned in **Data S5** with their putative Arabidopsis based annotation. The downregulated genes were also clustered into four major modules (**Figure S4**) that were annotated as photosystem related pathways (**Figure S5**).

### The clustered modules based on cumulative expression behaviour and their specific functional role in fiber development

Based on eigengene relationships, blockwise clustered modules were further categorized into two clades, clade I representing blue and turquoise module and clade II consists of red and yellow module with brown and green module outliers **(Figure 1f**). The expression profiling of genes belonging to clade I (blue and turquoise) showed higher expression during fiber elongation and SCW stages (**Figure 2a**). These two modules of clade I were highly connected to the hydroxyproline-rich glycoprotein family and Glycosyl hydrolases family protein respectively (**Figure 1f**, highlighted in pink color) which were acting as a central hub gene. GO analysis suggests that clade I were involved in lipid metabolism (**Figure 2d**) whereas Clade II shows their involvement in different organ development (**Figure 2d**) with higher expression level (especially for red and yellow modules) at fiber commitment stage. (**Figure 2b)**. Cytochrome P450 gene represented as a hub gene for yellow module, whereas red module containing genes were connected to the MADS-box family protein (**Figure 1f**, highlighted in pink color). The outlier module, green and brown modules also showed their relatively higher expression at the elongation and initiation stage (**Figure 2c**) with the central key players in the form of subtilase family gene and Glucose-methanol-choline oxidoreductase family protein respectively (**Figure 1f**, highlighted in pink color). Real-time expression analyses of hub genes validate the distinct expression profiling that representing the entire sets of genes for each module and reassure the involvement of modules for specific fiber developmental stages (**Figure 2e**).

**Figure 2.**
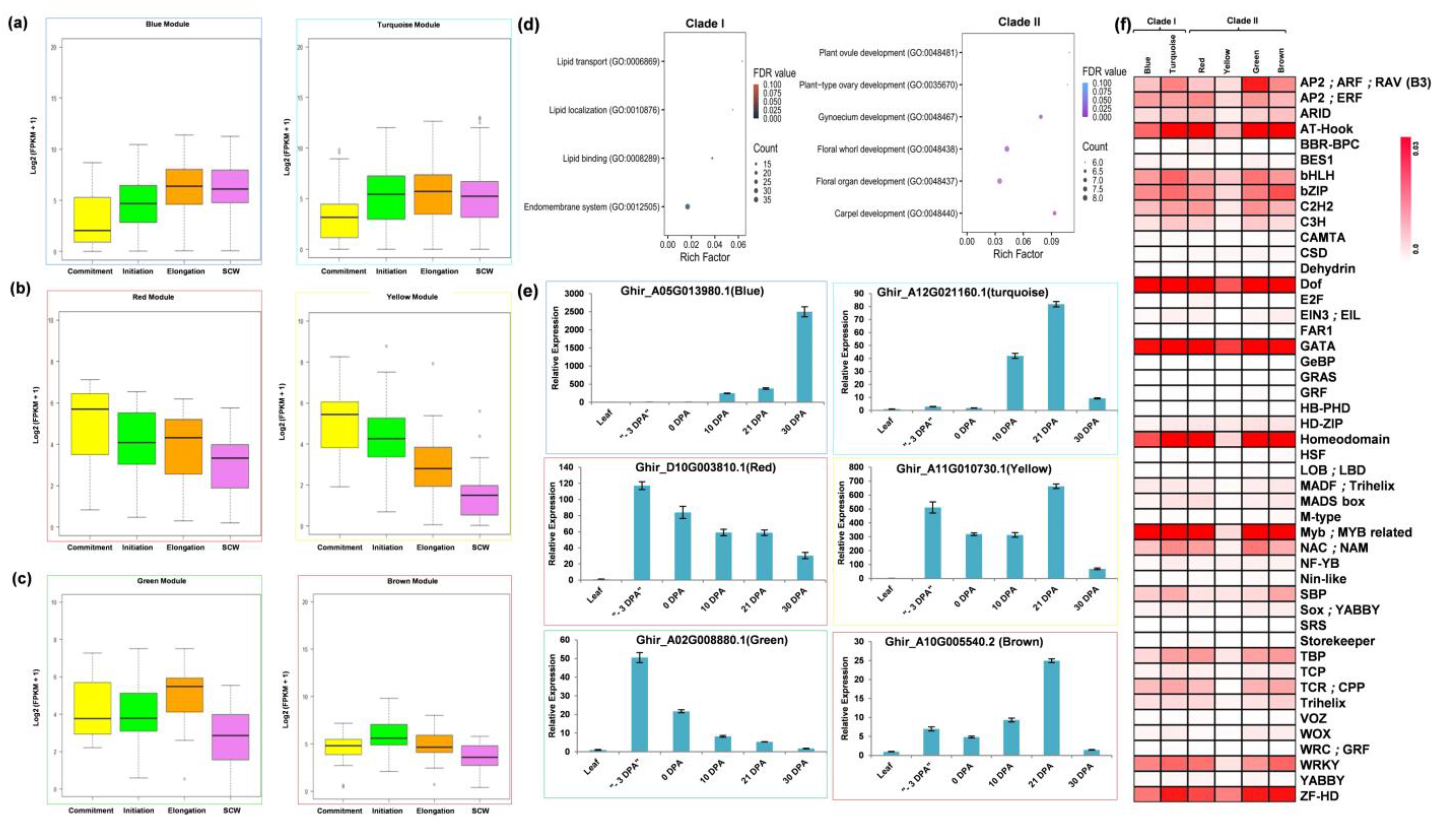
Functional characterizations of different modules in fiber tissue. Boxplot representation for expression profiling at Commitment, Initiation, Elongation and SCW fiber developmental stages for (a) Blue and Turquoise modules, (b) Red and Yellow modules and (c) Green and Brown modules. (d) Arabidopsis based GO enrichment of two major clades viz., Clade I and Clade II. The x-axis represents the ratio of total number of inputs to the background and y-axis shows the different GO Terms. The dot size represented the number of inputs for those particular terms while color code depicted the FDR value. (e) Real-time expression analysis of hub gene of all six modules. Hub genes are the representation of each module that confirmed their expression behaviour through real-time expression analyses in leaf tissues, −3 DPA, 0 DPA, 10 DPA, 21 DPA and 30 DPA of fiber developmental stages. (f) Binding frequency of cis-regulatory elements in the promoter region of genes present in all six modules viz., Blue, Turquoise, Red, Yellow, Green and Brown module. Color code ranges from 0 to 0.03, zero represents the lowest binding frequency of cis-regulatory element (CRE) while 0.03 depicted the higher binding frequency of CRE over the promoter regions.

### The enriched cis-regulatory elements and metabolic pathways of the conserved modules

We identified the binding frequency of the cis-regulatory element (CRE) in 1000 bp upstream of Transcriptional Start Site (TSS) using PlantPAN3.0 database (Chow et al., 2019). A total of 47 Transcription Factor (TFs) binding sites were identified in which Dof, GATA, ZF-HD TFs binding frequency were very prominent in all six modules (**Figure 2f) (Data S6)**. Homoedomain, MYB/Myb related, NAC and WRKY TFs binding sites frequency were significantly higher in all modules except for yellow. Four modules viz., turquoise, red, green and brown modules have high TATA Binding Protein (TBP) frequency while low in blue and yellow modules. Squamosa promoter binding protein (SBP) has a higher binding frequency in turquoise and brown module which was designated as an elongation specific module **(Figure 2a**,**2c)**.

Mapman pathway analysis revealed that the modules present in clade I were involved in cellular respiration and different type of metabolism pathways viz., carbohydrate, sucrose, starch and lipid metabolism **(Figure 3a**,**b)**. Thus, it was associated with cell wall formation through the upregulated expression of pectin and many glycoproteins **(Figure 3d)**. Many cytoskeleton related components like alpha, beta-tubulin and actin filament were also upregulated in clade I **(Figure 3c)** with oxidoreductase, transferases and hydrolases like enzymatic activities **(Figure 3h)**. Turquoise module of clade I was involved in chromatin modification **(Figure 3e)** and secondary metabolism pathway **(Figure 3g)** whereas blue module assists protein modification and phosphorylation via tyrosine kinase-like (TKL) superfamily upregulation **(Figure 3f)**. The upregulated genes of these two modules were involved in solute transport through the involvement of ABC, VHP PPase, MFS and TOC like superfamily transporter **(Figure 3i)**. Clade II modules show their involvement in solute transport through the upregulation of DMT and MFS superfamily **(Figure 3i)**. In the presence of extensin gene and LRR domain extensin (**Figure 3d**), yellow module involved in sucrose degradation and thus in cell wall formation **(Figure 3a)**. This module was mapped on auxin hormone pathway **(Figure 3j)** and belongs to MYB superfamily TFs **(Figure 3k)**. Other TFs like homoebox superfamily, MADS-box and trihelix were also aligned to this module whereas bZIP TFs were upregulated in green module **(Figure 3k)**. AP2/ERF and homeobox superfamily aligned to the turquoise module while no TFs were mapped on blue and brown module (**Figure 3k)**. Nevertheless, brown module’s genes facilitated the cuticle formation, phytosterol conjugation and pectin modification **(Figure 3d)** with an upregulated expression of ethylene hormone while red module manifested the cytokinin perception and signal transduction **(Figure 3j)**.

**Figure 3.**
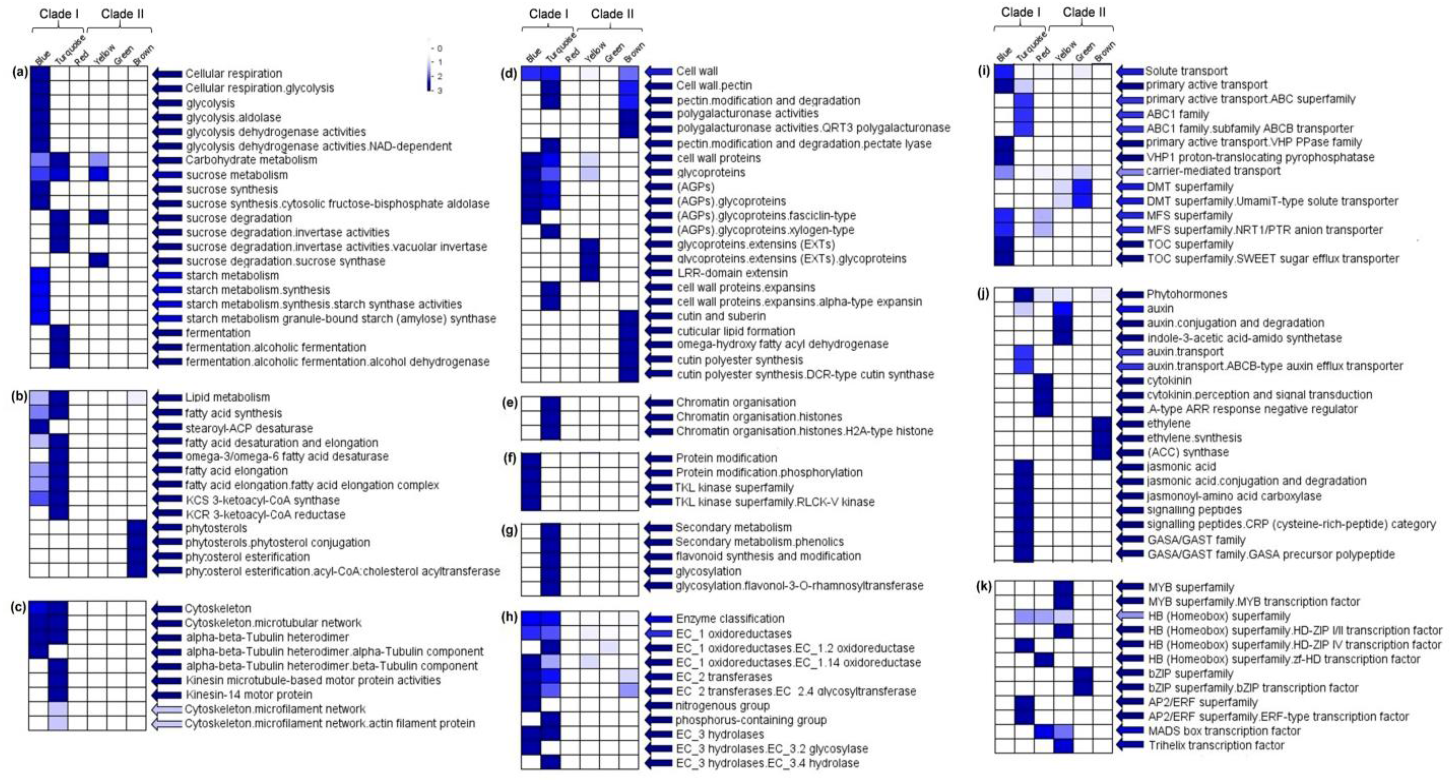
Mapman based significant functional classification of two different clades. These two clades consist of six modules that involved in different pathways (a) Cellular respiration and Carbohydrate metabolism (b) Lipid metabolism (c) Cytoskeleton molecules (d) Cell wall related functioning (e) Chromatin organization (f) Protein modification (g) Secondary metabolism pathway (g) Enzymatic action (i) Solute transportation (j) Phytohormone and (k) Transcription factors. BINs color code ranging from 0 to 3 with the vertical bar that represents the log2 expression value of each module.

### Transcriptional Biomarker: Initiation, Elongation and SCW specific features

A significant amount of RNAseq data of different fiber development stages (**Figure S6**) were used to identify the unique transcriptional biomarkers (Riedmaier and Pfaffl, 2013). After the normalization, the data of initiation, elongation and SCW stages was clustered in two panels based on their expression similarities. Data from the initiation stage showed a distinct cluster while that of elongation and SCW showed overlap (**Figure S7**). Thus, we defined the initiation and elongation or SCW specific transcriptional biomarker based on informative features of high coverage transcriptome data (**Figure S8**). The top 20 important scoring genes (RReliefF score) were plotted with their z score value (**Figure 4a**). Positive z score value of alpha/beta hydrolases superfamily protein, IQ domain26, aquaporin transporter, gibberellin regulated protein and peroxidase gene family at initiation stages indicate them as a transcriptional biomarker for fiber initiation. Similarly, several genes showed higher z score value at the elongation or SCW stage viz., HAD like superfamily, phospholipases A2A protein, prolin transporter 1, SWEET sugar transporter and some predicted protein (Ghir_A05G0105660, Ghir_A06G004130, Ghir_D09G009930, Ghir_D07G020490). Thus these genes were considered as a transcription biomarker for elongation or SCW stages. The relative expression analysis using quantitative real-time PCR (qRTPCR) strongly reaffirms the expression profiling of identified transcriptional biomarkers (**Figure 4b**,**c**) where we found the initiation, elongation or SCW specific features.

**Figure 4.**
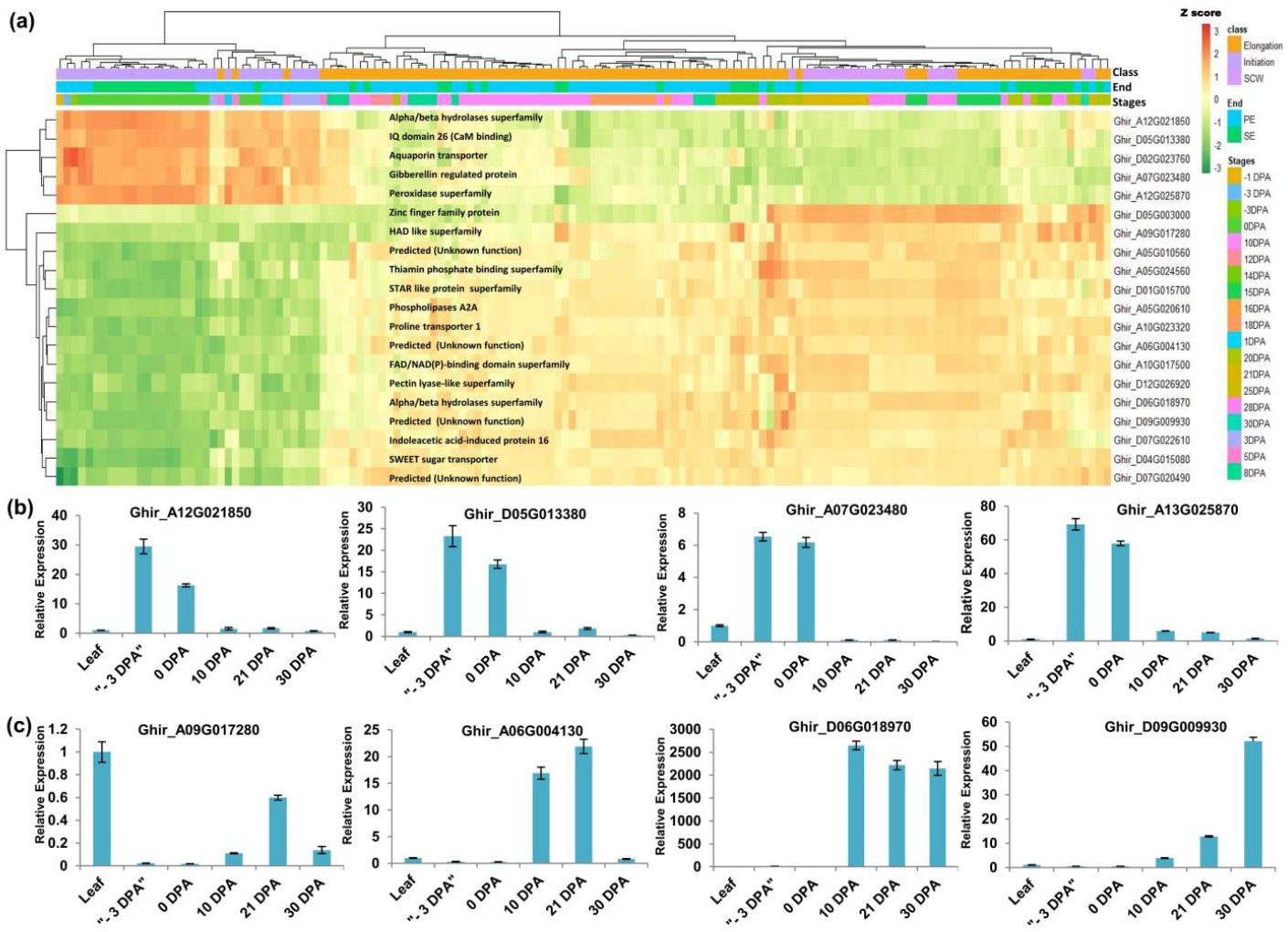
Interpretation of Transcriptional biomarker for Initiation and Elongation or SCW specific stages. (a) Expression matrix of top twenty high RRelif scoring genes among the different DPA stages. The normalized z score value was represented in the form of an expression matrix. The green color represents the lowest expression value while the orange color represented the highest expression value. The putative function of each gene was mentioned along each row of genes. Relative expression pattern of randomly selected four (b) Initiation specific transcriptional biomarker and (c) Elongation or SCW specific transcriptional biomarker in leaf tissue and different stages of fiber tissues viz., −3 DPA, 0 DPA, 10 DPA, 21 DPA and 30 DPA.

### The exclusively expressed gene variants in cotton fiber development

Besides 302 up-regulated genes in OF, we identified 20 transcripts (representing gene variants) that expressed exclusively in OF as compared to LRS. Expression of these transcripts was very significant as we processed approximately ∼400X uniquely mapped read coverage to the genome. Although, the average expression level of these transcripts (0.99) was low in comparison to the average expression level of earlier identified 302 upregulated genes (5.78). It is very interesting to note that the expression of these gene variants was very specific (**Table 1**) to the OF tissues as compared to its other variants which showed somewhat higher expression in non-fiber tissue (LRS). Arabidopsis based annotation analysis revealed that these gene variants belong to ELO gene, PDF2, WRKY and bHLH TFs, several enzymes (glycerol-3 phosphatase dehydrogenase, benzoquinone reductase) and many hypothetical proteins. The qRTPCR analyses represent the typical expression as determined by *In-Silico* transcriptome analysis. In real-time expression analysis, hypothetical protein (Ghir_D10G025770.3) expressed at −3 DPA (Day Post Anthesis) and 0 DPA and serine threonine-protein kinase transcript (Ghir_D05G003410.3) shows significant expression at the elongation stage (**Figure 5a**) with null expression value in leaf tissue.

**Table 1.**
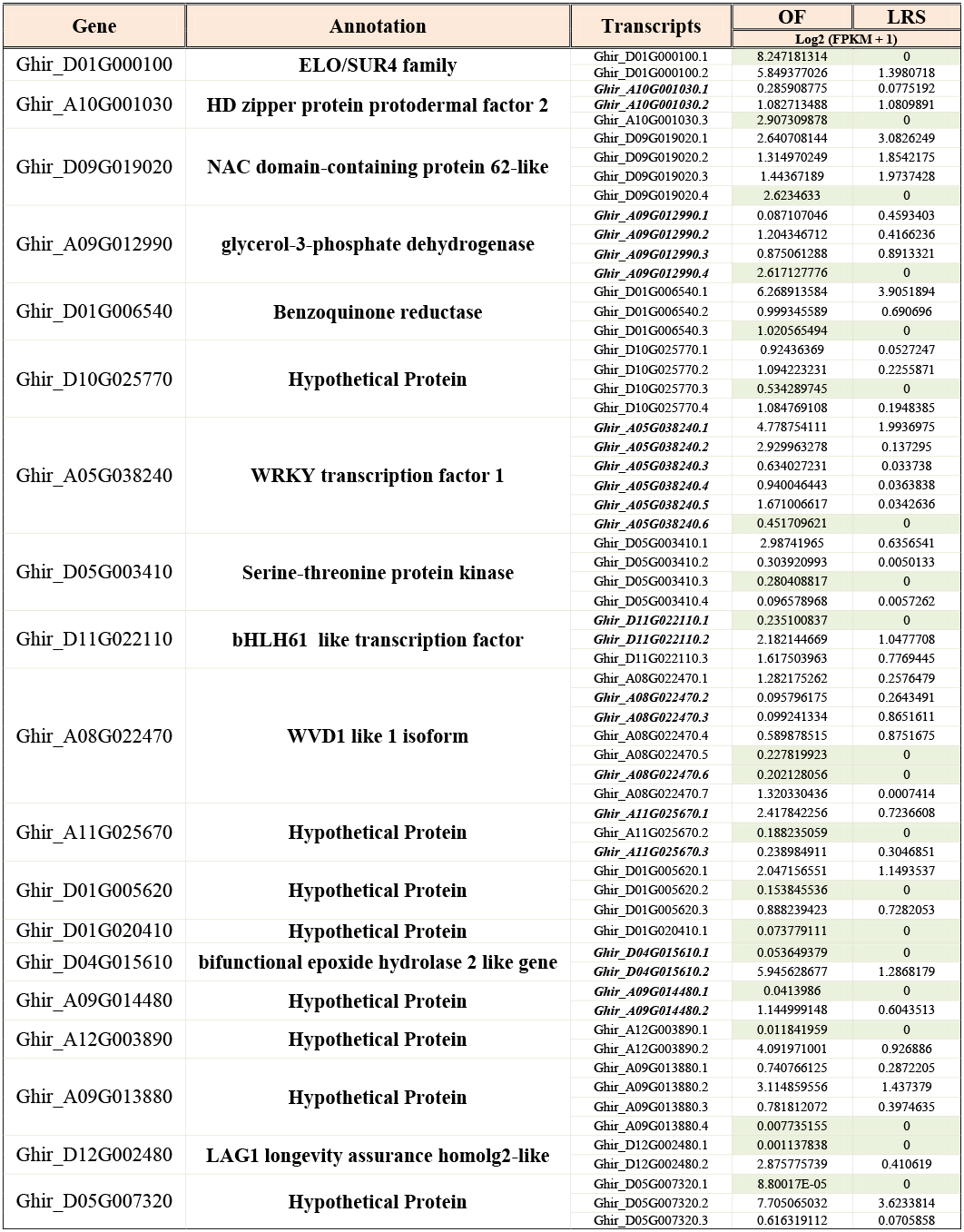
Detail information of Exclusively Expressed Transcripts (EETs) including their Arabidopsis based annotation and log2(FPKM +1) value in OF and LRS tissues. Identified EETs were highlighted with olive green colored cell whose expression was null in LRS tissue. Some identified transcripts have identical nucleotide sequences which were represented by the Bold and Italic font in the transcripts column.

**Figure 5.**
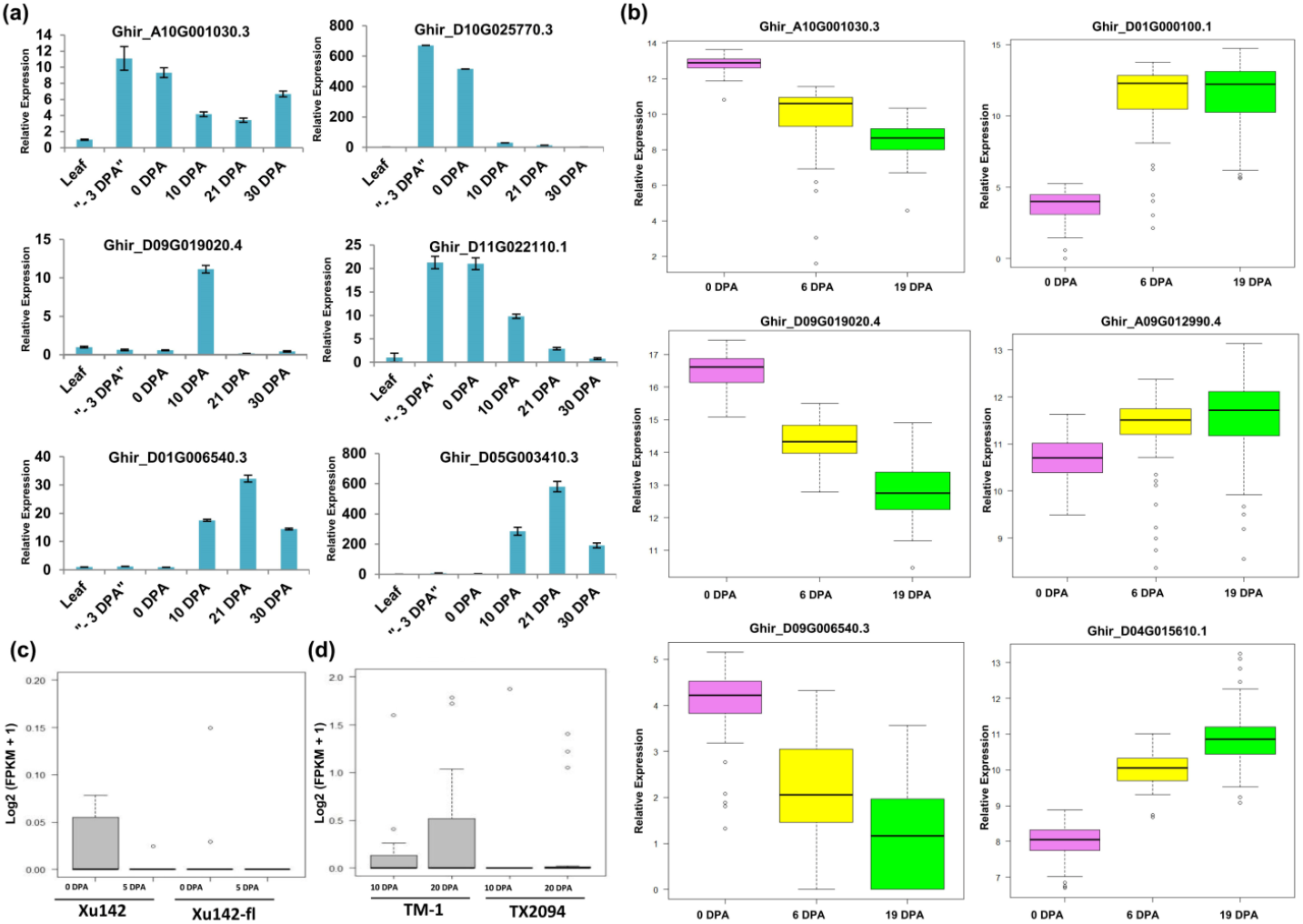
Validations of Exclusively Expressed Transcripts (EETs). (a) Real-time expression validation in leaf and different stages of fiber tissue viz, −3 DPA, 0 DPA, 10 DPA, 21 DPA and 30 DPA for six selected EETs viz., top five highly expressed transcripts and one low expressed transcript. *In-silico* expression pattern of EETs (b) for domestication process in modern-day cotton (™-1) and wild cotton (TX2094) at 10 and 20 fiber developmental stages and (c) for fiber specific pattern in wild cotton (Xu142) and *fuzzless* natural mutant (Xu142*fl*) at 0 DPA and 5 DPA. (d) Relative expression of Nanostring nCounter gene expression assay of six selected transcripts in 100 genotypes at 0 DPA, 6 DPA and 19 DPA fiber developmental stages in 2018 growing seasons at Hyderabad city of India.

### Exclusively Expressed Transcripts are positively selected during evolution and domestication

Exclusive Expressed Transcripts were very specific to the fiber structures so to substantiate their importance in cotton genome, we checked the collective expression of these transcripts in fiberless mutant, Xu142*fl* and wild cotton cultivar, Xu142 (**Figure 5b**). In fiberless mutant cotton, these transcripts show null expression (Fragments Per Kilobase Million (FPKM) value = zero) at 0 DPA and 5 DPA fiber developmental stages, indicating their potential role in various stages of fiber development, especially at the initiation stage. We also checked the expression profiling of these 20 gene variants during the domestication process by comparing the wild cotton variety, TX2094 and domesticated cotton variety, ™-1 (**Figure 5c**). The cumulative expression of 20 EETs was significantly low at 10 DPA and 20 DPA in wild variety than the domesticated variety. These observations further corroborating the functional importance of these genes in the cotton fiber developmental stages during the different evolutionary processes.

### The expression of EETs correlates to the quality parameters of cotton fiber

We selected five highly expressed transcripts (log2(FPKM +1) >1) viz., ELO/SUR4 family, PDF2, NAC domain-containing protein 62 like, glycerol-3-phosphate dehydrogenase, benzoquinone reductase and one low expressed transcript (log2(FPKM + 1) < 0.1) bifunctional epoxide hydrolase 2 for expression and correlation study in 100 genotypes (**Data S7**). Different fiber quality and fiber-yield traits were estimated for two consecutive years viz; 2018 at Hyderabad, 2019 at Aurangabad and Hyderabad. Aurangabad and Hyderabad are two different cities of Maharashtra and Telangana state, respectively which reported as the leading state in cotton acreage in India (Nagrale et al., 2020). The phenotypic relevance of these fiber quality traits was highly correlated with each other in two different seasons (2018 and 2019) (**Figure S9**) and different locations (Hyderabad and Aurangabad) (**Figure S10**). Expression of selected exclusively expressed transcripts was further examined using nanostring nCounter technology at three fiber developmental stages viz., 0 DPA, 6 DPA and 19 DPA in 100 genotypes of cotton that were grown in 2018 at Hyderabad city of India (**Figure 5d**). The nCounter expression analysis confirmed that three transcripts viz., ELO/SUR4, glycerol-3-phosphate dehydrogenase and bifunctional epoxide hydrolase 2 like gene were significantly highly expressed at 6 DPA and 19 DPA in comparison to 0 DPA while three other transcripts PDF2, NAC domain-containing protein 62 like and benzoquinone reductase were highly expressed at 0 DPA. The expression of exclusive transcripts observed in nCounter assay confirmed their expression pattern which was obtained through *In-Silico* analysis and thus further validating our results.

Pearson correlation coefficient and the statistical significance test were used to demonstrate the linear relationship between the normalized nanostring nCounter data and fiber quality traits of 100 genotypes. In correlation and statistical analysis, ELO/SUR4 family gene variant (Ghir_D01G000100.1) exhibits a significant positive correlation with fiber strength, micronaire, boll number and cotton yield at 0 DPA (**Table 2**). This transcript also correlated with elongation, micronaire and boll weight traits at 6 DPA and only with the boll weight at 19 DPA. An unlikely significant negative correlation was seen with the micronaire at 6 DPA in 2019 at Hyderabad city. NAC domain-containing protein 62 like (Ghir_D09G019020.4) exhibits a negative significant correlation with micronaire trait at 0 DPA and 6 DPA while the positive correlation with fiber length at 6 DPA in two consecutive years (2018, 2019) as well as in two different locations (Hyderabad, Aurangabad). Contrasting behavior of the NAC domain gene was observed in 2018_H and 2019_A data at 0 DPA where we observed a significant negative and positive correlation respectively with fiber elongation trait. At 0 DPA, HD zipper protein protodermal factor 2 transcripts (Ghir_A10G001030.3) showed its significant positive correlation with the fiber length, uniformity index and fiber strength while significant negative correlation with boll weight at 6 DPA for 2019_H (**Table 2**). Glycerol-3-phosphate dehydrogenase (Ghir_A09G012990.4) and bifunctional epoxide hydrolase 2 like (Ghir_D04G015610.1) transcripts showed a significant positive and negative correlation with elongation and micronaire respectively at 0 DPA. The Bifunctional epoxide hydrolase 2 like transcript was also positively correlated to the fiber elongation at 0 DPA and 6 DPA and cotton yield at 6 DPA. The benzoquinone reductase transcript (Ghir_D01G006540.3) displayed a significant negative correlation with the micronaire at 0 DPA, fiber length and boll weight at 6 DPA and fiber length, uniformity index, fiber strength, elongation, boll number and cotton yield at 19 DPA (**Table 2**). All selected transcripts show the significant negative correlation with micronaire at 0 DPA and 6 DPA, exceptional ELO at 0 DPA and benzoquinone reductase at 6 DPA. None of the transcripts give a significant correlation at 19 DPA for micronaire trait. All the mentioned correlations were significantly affected the fiber-related traits either positively or negatively.

**Table 2.** Functional validation of EETs for different fiber quality traits and fiber yield-related traits. Pearson Correlation Coefficient of Nanostring nCounter gene expression value of 100 genotypes in 2018 year at Hyderabad city with fiber-related traits in three different conditions viz., 2018_H, 2019_H and 2019_A. 2018 and 2019 represented the two consecutive growing seasons while H and A depicted as Hyderabad and Aurangabad city of India, respectively. Green colored cell represented the positive correlation whereas red colored cell represented the negative correlation value. Significant correlation were denoted with the asterisk mask (*) in which *, **, *** used for the p-value ≤ 0.05, ≤ 0.01, ≤ 0.005, respectively.

| Stages | Gene | Annotation | Fiber Quality Traits |  |  |  |  |  |  |  |  |  |  |  |  |  |  | Fiber Yield Traits |  |  |  |  |  |
| --- | --- | --- | --- | --- | --- | --- | --- | --- | --- | --- | --- | --- | --- | --- | --- | --- | --- | --- | --- | --- | --- | --- | --- |
|  |  |  | Fiber Length |  |  | Uniformity Index |  |  | Fiber Strength |  |  | Elongation |  |  | Micronaire |  |  | Ball Number |  | Ball Weight |  | Seed Cotton Yield |  |
|  |  |  | 2018_H | 2019_H | 2019_A | 2018_H | 2019_H | 2019_A | 2018_H | 2019_H | 2019_A | 2018_H | 2019_H | 2019_A | 2018_H | 2019_H | 2019_A | 2019_H | 2019_A | 2019_H | 2019_A | 2019_H | 2019_A |
| 0DPA | Ghir_D01G000100.1 | ELO/SUR4 family | 0.037 | 0.18 | 0.21 | 0.018 | 0.054 | 0.16 | 0.049 | 0.17 | 0.29** | 0.054 | 0.048 | -0.16 | 0.22* | 0.29** | 0.23* | -0.044 | 0.3** | 0.19 | 0.0004 | -0.034 | 0.33** |
|  | Ghir_D09G019020.4 | NAC 62-like | -0.012 | 0.049 | 0.001 | -0.062 | 0.16 | 0.12 | -0.14 | 0.015 | -0.04 | -0.23* | 0.0059 | 0.56*** | -0.26* | -0.39*** | -0.13 | 0.021 | -0.28* | -0.043 | 0.23* | 0.031 | -0.16 |
|  | Ghir_A10G001030.3 | PDF2 | 0.27* | 0.27* | 0.3** | 0.21 | 0.12 | 0.3** | 0.17 | 0.16 | 0.22* | 0.03 | -0.023 | 0.071 | -0.1 | -0.12 | -0.11 | 0.15 | 0.063 | 0.1 | 0.039 | 0.12 | 0.011 |
|  | Ghir_A09G012990.4 | glycerol-3-phosphate dehydrogenase | 0.024 | 0.036 | 0.011 | 0.11 | 0.2 | 0.18 | 0.011 | 0.12 | 0.001 | -0.11 | 0.1 | 0.59*** | -0.12 | -0.24* | -0.004 | 0.16 | -0.064 | 0.00004 | 0.19 | 0.36*** | -0.009 |
|  | Ghir_D01G006540.3 | Benzoquinone reductase | 0.18 | 0.13 | 0.16 | 0.034 | 0.14 | 0.14 | -0.0005 | 0.046 | 0.007 | -0.12 | 0.089 | 0.15 | -0.27* | -0.25* | -0.16 | -0.21 | -0.037 | -0.075 | 0.14 | 0.045 | 0.025 |
|  | Ghir_D04G015610.1 | bifunctional epoxide hydrolase 2 like | -0.037 | 0.043 | -0.005 | -0.035 | 0.1 | 0.074 | -0.14 | 0.07 | -0.11 | -0.082 | -0.015 | 0.51*** | -0.16 | -0.29** | -0.001 | -0.011 | -0.19 | -0.045 | 0.16 | 0.041 | -0.16 |
| 6DPA | Ghir_D01G000100.1 | ELO/SUR4 family | -0.064 | -0.1 | -0.14 | 0.031 | -0.14 | 0.014 | -0.098 | -0.11 | -0.13 | -0.023 | -0.043 | 0.25* | -0.019 | -0.25* | -0.1 | -0.17 | -0.097 | -0.18 | 0.25* | -0.011 | -0.048 |
|  | Ghir_D09G019020.4 | NAC 62-like | 0.22* | 0.31** | 0.23* | 0.086 | 0.18 | 0.15 | 0.068 | 0.2 | 0.14 | -0.024 | 0.1 | -0.002 | -0.2 | -0.23* | -0.1 | 0.11 | 0.14 | 0.081 | -0.052 | 0.031 | 0.11 |
|  | Ghir_A10G001030.3 | PDF2 | 0.03 | -0.08 | -0.055 | 0.052 | -0.065 | -0.022 | -0.085 | -0.072 | -0.16 | -0.023 | -0.012 | 0.076 | 0.015 | -0.19 | -0.17 | -0.078 | -0.078 | -0.26* | 0.22 | -0.018 | -0.074 |
|  | Ghir_A09G012990.4 | glycerol-3-phosphate dehydrogenase | 0.12 | 0.035 | 0.017 | 0.21 | 0.039 | 0.1 | 0.0064 | 0.017 | -0.059 | -0.011 | -0.017 | 0.12 | -0.015 | -0.28* | -0.2 | -0.068 | -0.046 | -0.13 | 0.33** | 0.1 | 0.013 |
|  | Ghir_D01G006540.3 | Benzoquinone reductase | -0.066 | -0.26* | -0.17 | -0.17 | -0.19 | -0.093 | -0.14 | -0.18 | -0.064 | 0.0002 | -0.18 | -0.19 | -0.19 | 0.24* | 0.059 | 0.0083 | 0.11 | -0.12 | -0.27* | -0.034 | -0.18 |
|  | Ghir_D04G015610.1 | bifunctional epoxide hydrolase 2 like | 0.1 | 0.12 | 0.033 | 0.089 | 0.022 | 0.046 | 0.074 | 0.14 | -0.022 | 0.26* | -0.044 | 0.15 | -0.16 | -0.23* | -0.11 | 0.099 | 0.028 | -0.056 | 0.2 | 0.25* | 0.084 |
| 19DPA | Ghir_D01G000100.1 | ELO/SUR4 family | 0.0022 | 0.1 | 0.028 | 0.046 | 0.11 | 0.12 | 0.019 | 0.089 | 0.063 | 0.043 | 0.11 | 0.16 | 0.011 | 0.062 | 0.065 | 0.1 | -0.055 | 0.26* | 0.036 | 0.099 | 0.025 |
|  | Ghir_D09G019020.4 | NAC 62-like | 0.062 | 0.11 | -0.05 | 0.086 | 0.071 | -0.014 | 0.066 | 0.11 | -0.05 | -0.066 | 0.15 | 0.11 | -0.001 | 0.032 | 0.038 | -0.12 | -0.2 | 0.072 | -0.043 | 0.019 | -0.12 |
|  | Ghir_A10G001030.3 | PDF2 | -0.067 | -0.072 | 0.042 | -0.07 | -0.12 | -0.014 | -0.17 | -0.04 | -0.012 | -0.03 | -0.24* | 0.058 | -0.13 | -0.12 | -0.054 | 0.023 | 0.046 | -0.13 | -0.06 | 0.031 | 0.042 |
|  | Ghir_A09G012990.4 | glycerol-3-phosphate dehydrogenase | 0.064 | 0.11 | 0.13 | 0.051 | 0.11 | 0.099 | 0.042 | 0.12 | 0.15 | -0.0073 | 0.1 | -0.0026 | 0.068 | 0.075 | 0.093 | 0.1 | 0.1 | 0.16 | -0.058 | 0.2 | 0.15 |
|  | Ghir_D01G006540.3 | Benzoquinone reductase | -0.21 | -0.25* | -0.28* | -0.089 | -0.26* | -0.23* | -0.14 | -0.27* | -0.2 | -0.086 | -0.27* | -0.11 | 0.052 | -0.028 | 0.03 | -0.27* | -0.091 | -0.13 | 0.024 | -0.23* | -0.016 |
|  | Ghir_D04G015610.1 | bifunctional epoxide hydrolase 2 like | -0.037 | 0.0047 | 0.058 | -0.064 | 0.012 | 0.0061 | -0.059 | 0.036 | 0.066 | -0.11 | -0.046 | 0.09 | -0.12 | -0.12 | -0.063 | 0.019 | 0.0047 | -0.0056 | -0.05 | 0.15 | 0.03 |

## Discussion

Cotton genome was refined over the year by different independent groups to get the well-annotated genes and unravel the evolution dynamics for fiber-specific features (Yang et al., 2020). Additionally, to understand the molecular and biological functions of genes involved in cotton fiber growth and development, the multi-omics approach was also taken into consideration (Pang et al., 2009; Rai et al., 2013; Song et al., 2015; Zou et al., 2016; Song et al., 2017; Kumar et al., 2018; Ayubov et al., 2019; Li et al., 2019). In recent years, many researchers were reannotated the cotton genome sequence by third and fourth generation sequencing techniques to get the high contiguity genome (Pan et al., 2020). Prior to cotton genome sequence was available, several transcriptome data (Wang et al., 2010; Nigam et al., 2014; Peng et al., 2014; Cheng et al., 2016; Parekh et al., 2018) were generated that would not have been mapped properly due to the unavailability of a well furnished reference genome. Recent efforts have been carried out for the identification of important molecular signatures for fiber-related traits, in which around five to seven fold coverage transcriptome data were generated and mapped on the draft genome of cotton (Fang et al., 2017; Ma et al., 2018). The low coverage on draft genome may not be used for the identification of low expressed gene and their variants that might be crucial for fiber development. Similarly in an eQTL identification study, only 10 billion reads were used for 251 cotton accessions (Li et al., 2020). This study was only focused on the initiation of secondary cell wall synthesis at 15 DPA. Till date, no studies are available that used high coverage cotton transcriptome data on new assembled genome (Hu et al., 2019; Wang et al., 2019; Yang et al., 2019; Chen et al., 2020; Huang et al., 2020) for all fiber developmental stages. In our current study, we elucidated many fiber specific features for different fiber developmental stages categorizing into commitment, initiation, elongation and secondary cell wall synthesis stages where we processed and reanalyzed the 356 publicly available cotton transcriptome data. These large-scale data were comprised of 18 different tissues and 34 cultivars (**Figure 1A, Figure S1**) that span ∼400 fold coverage to the newly assembled genome (Wang et al., 2019). The high coverage data helped us in the identification of important fiber features that were not reported yet for different fiber developmental stages which subsequently correlated with the different fiber-related traits.

The comprehensive clustering of fiber-specific genes based on their expression resulted in six distinct modules, which are further grouped into two prominent clades to represent the stage-specific expression (**Figure 1e-f, 2a-c**). Clade-II significantly represents commitment-specific modules namely, yellow and red modules which were involved in different organ development while clade I modules viz., blue and turquoise were involved in the fiber cell elongation and maturation through lipid metabolism processes (**Figure 2d**). The yellow module’s genes were annotated as MYB, HD-ZIP, MADS, and Trihelix transcription factors **(Figure 3k) (Figure S4)**. The role of these transcription factors like MYB/ MYB like (Walford et al., 2011) (MacHado et al., 2009) and homeodomain leucine zipper gene (Walford et al., 2012) in the fiber initials further affirmed the specificity of this module for commitment specific expression. Interestingly, although the expression of cell fate-determining factors - MYB and HD-ZIP is significantly high in the yellow module however promoter analysis indicates the poor enrichment of their binding site in the promoter of the same module (**Figure 2f**). This indicates that expression of MYB and HD-ZIP are governed by different regulators at the commitment stage but they themselves regulate the expression of genes of other modules as represented through CRE frequency in the promoter regions (**Figure 2f**). Further, the significance of the yellow module is substantiated by the presence of genes belonging to the auxin and indole-3-acetic acid amido synthase pathway whose role was well established for cell fate determination (Zhang et al., 2011; Xu et al., 2019). The role of auxin efflux and influx is well documented in the fiber initials at −1 DPA (Petrášek et al., 2006; Zhang et al., 2017) and fiber protrudes through the upregulation of genes belonging to extensin related glycoproteins. We also observed significant expression of extensin related proteins and LRR domain-containing proteins at −3 to 0 DPA stages in yellow module for the fiber development (**Figure 3d**). Another module of clade I viz., red module shows the upregulation of the cytokinin and cytokinin perception and signal transduction genes including A type ARR negative regulator **(Figure 3j)**. Thus, fiber development may follow a cascade where the antagonistic behavior of auxin and cytokinin (Zeng et al., 2019) has been maintained by the upregulation of A-type ARRs. The functional significance of genes identified in each module was also ascertained since many genes fall in the genic and intergenic region that were associated with several fiber-related traits as shown in previously identified genome-wide association study (Ma et al., 2018). All modules demonstrated their association with different fiber quality and fiber yield-related traits in which yellow module exhibit the maximum number (6 out of 27) of association with different traits (**Table S1**).

Apart from the conventional studies, genome-wide transcriptional data of different fiber developmental stages facilitated the identification of transcriptional biomarkers for different stages of development. For cotton breeders, molecular markers are useful to identify the genomic regions to underpin the economic traits (Malik et al., 2014). We thus identified fiber initiation and elongation or SCW specific transcriptional biomarkers whose expression was typical for those stages. We identified gibberellin-regulated protein (Ghir_A07G023480) as a biomarker for initiation stage (**Figure 4a**,**b**). The increased level of gibberellic acid (Tu et al., 2007; Xiao et al., 2010) was reported during the early stages of fiber development which might regulate gibberellin regulated protein. IQ domain 26, a calmodulin-binding domain-related family protein was also identified as biomarker for ™3 DPA and 3 DPA (**Figure 4b**) which is consistent with previous reports, suggesting its role in calmodulin signaling pathway at fiber initials stage (Taliercio and Boykin, 2007). According to the previous study, the aquaporin transporter was responsible for generating the turgor pressure in fiber cells at 3 DPA to 20 DPA through intense water influx (Li et al., 2013). In our analysis, a member of aquaporin transporter (Ghir_D02G023760) was identified as a transcriptional biomarker (**Figure 4a**,**b**) for initiation stage that might be facilitating the water influx in fiber initials. In cotton, peroxidase superfamily maintained the hydrogen peroxide (H2O2) homeostasis for fiber initiation and elongation (Guo et al., 2016). Peroxidases are superfamily protein which exhibits their diverse role in different fiber developmental stages whereas, in our study, one of its members (Ghir_A12G025870) appeared as a transcriptional biomarker for initiation stages (**Figure 4a**,**b)**. Likewise, Ghir_A12G021850 and Ghir_D06G018970 are the two members of alpha beta hydrolases was identified as a biomarker for initiation as well as for elongation or SCW stages respectively (**Figure 4a**,**b**,**c**), suggesting the importance of superfamily protein in cotton fiber development. The expression levels of all identified transcriptional biomarkers are very specific to their stages that authenticate our findings. Through this study, many predicted proteins (Ghir_A06G004130, Ghir_D09G009930) were emerged out as important features **(Figure 4b)** that need to be investigated further to understand their role in fiber development.

High coverage transcriptome data were used to identify the full sequence diversity of the genome, including lowly expressed transcripts (Fu et al., 2014). In the present study, approximately six billion uniquely mapped reads were used to discover the new genes or transcripts for fiber-specific expression. Our study identified 20 EETs that were exclusively expressed in fiber tissues. The importance of exclusivity of these transcripts can be seen by their higher expression in OF in comparison to null expression in LRS tissue **(Figure 5a)** (**Table 1**), indicating their importance in fiber development. During the domestication of cultivated cotton species, these transcripts might be positively selected in modern cotton for fiber specific features as we observed almost nil expression in the wild type cotton (TX2094) (**Figure 5b**) as well as in fiberless mutant (Xu142*fl*) (**Figure 5c**). The comprehensive expression analysis of EETs using nCounter gene expression assay involving 100 genotypes and three development stages confirmed the different stage-specific expression of EETs (**Figure 5d**). The correlation analysis established a significant correlation between the expression of EETs and different fiber quality and yield-related traits (**Table 2**). The significant correlations further ascertain the importance of EETs in cotton fiber development identified in our study.

One of the tested EET, PROTODERMAL FACTOR 2 (Ghir_A10G001030.3) was identified to be highly expressed at 0 DPA (**Figure 5d**) and significantly correlated to the fiber length (**Table 2**). This was consistent with the previous report wherein the role of PROTODERMAL FACTOR 1 and its mechanism in fiber development was well documented (Deng et al., 2012). Yet another EET belongs to NAC family gene, which was known previously for its involvement in fiber development (Shang et al., 2016). Our correlation study reveals that NAC negatively correlates to micronaire at 0 DPA and positively correlates with fiber length at 6 DPA (**Table 2**). This study reaffirms the previous studies where NAC has been documented for fiber development both positively (Zhang et al., 2018) and negatively (Sun et al., 2020). ELO transcript (Ghir_D01G000100.1) also correlates positively with the micronaire at 0 DPA. ELO is a member of the elongase gene family that putatively involved in the synthesis of very long chain fatty acids (VLCFs) (Qin et al., 2007a; Zhang et al., 2015) which ultimately acts as precursors for the sphingolipid biosynthesis to promote cotton fiber elongation (Qin et al., 2007a; Qin et al., 2007b). Our nCounter gene expression suggests that ELO’s expression initiated early at 0 DPA, thus it is very interesting to validate further that how the early expression of ELO influences micronaire trait. Our correlation study established an inverse relationship between fiber elongation and micronaire, which may be crucial for sustain the yarn strength (Mathangadeera et al., 2020) by combining properties of fiber length, density and maturity (Long et al., 2013).

Our extensive and comprehensive RNAseq data thus resulted into the identification of several new genes whose expression might be crucial for fiber development. Many of these newly identified genes may influence important agronomical cotton fiber traits and thus our expression and correlation analysis involving 100 genotypes and three developmental stages further substantiate the importance of identified genes.

## Methods

### Differential Quantification and Cluster Analysis of Cotton transcriptome data

More than 350 RNAseq datasets of the mock-treated or untreated cotton samples were collected from the NCBI SRA database that was available till September 2018. These datasets are of mixed type (paired-end and single-end) with 175 bp average length, comprised of 18 tissues and 34 cultivars. SRA data were converted into fastq file and processed the quality control followed by the adaptor trimming by the Trimmomatic tool (Bolger et al., 2014) wherever required. Eight billion filtered reads were mapped on the *Gossypium hirsutum* reference genome (Wang et al., 2019) through the STAR aligner (Dobin et al., 2013) with default settings.

Five tissues: Ovule, Fiber, Leaf, Root, and Seed have a higher number of RNAseq samples were selected for downstream analysis with the minimum 60 % uniquely mapped reads that span the 400X coverage to ∼2.8 Gb cotton genome size. 314 samples of selected five tissues were grouped into two major tissues; OF (Ovule + Fiber) and LRS (Leaf + Root+Seed) for fiber specific differential quantification using the Ballgown R package (Pertea et al., 2016). After the removal of low abundance genes across the samples, 973 genes were differentially expressed with FDR < 0.005 and fold change > 2. Among the differentially expressed genes (DEGs), 302 upregulated and 671 downregulated genes were subjected to the Weighted Gene Co-expression Network Analysis (Langfelder and Horvath, 2008) that categorized the DEGs into different modules. Blockwise modules were used with the soft threshold power of 10 to generate the module wise clustering where the minimum module size was set to 10 with unsigned Topological Overlap Matrix. Clustering was based on co-expression behaviour among the genes. By defining the degree of centrality between gene, hub gene was also identified for these modules which were eventually visualized through Cytoscape plugin (Paul Shannon et al., 1971).

### Consolidation of different DPA into four overlapping stages

To check the expression profiling of different modules in fiber development stages, we consolidate the samples in four major subsets according to the different DPA viz.: ‘-3 DPA’, ‘-1 DPA’, ‘0 DPA’ into commitment, ‘1 DPA’, ‘3 DPA’ were merged into initiation stages, ‘5 DPA’, ‘8 DPA,’ ‘10 DPA’, ‘12 DPA’,’14 DPA’, ‘15-16DPA’,’18 DPA’,’20 DPA’ grouped for the elongation, while ‘21 DPA’ ‘25-28 DPA’,’30 DPA’,’35 DPA’ and ‘40 DPA’ were defined for SCW stages. All four subsets were assumed to be representative of fiber overlapping development stages. Further, we quantitate the transcripts through the StringTie assembler (Pertea et al., 2016) for each stage. These subsets were used to check the *In-silico* expression pattern for downstream analysis.

### Functional enrichment and cis-regulatory elements identification of differently clustered modules

Gene Ontology analysis of differentially regulated genes was carried out through the AgriGO database (Du et al., 2010). For the GO enrichment, arabidopsis genome was taken as a reference. Significantly enriched, no-redundant GO Terms were visualized through R package (Team, 2013). Mapman analysis was carried out where data points in each bin were averaged and resulting p-value were adjusted to the Bonferroni.

To check the regulatory elements in all six modules, promoter analysis was conducted through the PlantPAN3.0 database (Chow et al., 2019). We retrieve the genomic sequences from 1000bp upstream of TSS of each gene in all modules and scanned further for identification of CRE binding site. The frequency binding site for CRE for each module was calculated as – number of TF binding sites in the promoter sequences of each module multiplied by the motif length which is further divided by the total number of genes present in that module multiplied by 100.

### Identification and *In-Silico* validation *of* exclusively expressed transcripts in fiber tissue

We validated the expression of 20 identified EETs in the mutant line as well as in wild cotton. For *In-Silico* validation, we selected different SRA data, other than the previous processed SRA for Xu142*fl* mutant line and TX2094 domesticated cotton cultivar. We accessed the SRA049496 data and mapped the filtered reads on the reference genome (**Data S8**) to quantitate the exclusively expressed transcripts level at 0 DPA and 5 DPA in the natural fiberless mutant. Additionally, to scrutinize the expression status of exclusively expressed transcripts during the domestication process, we retrieve the RNAseq data of ™-1, domesticated cotton and TX2094, wild cotton cultivar on 10 DPA and 20 DPA fiber samples of SRP017061 accessions. All the accessed SRA data were processed in a similar way by which the previously downloaded SRA data were processed for quantification of OF and LRS tissues.

### Data mining for Initiation, Elongation and SCW Specific Transcriptional Biomarker

We thoroughly pre-processed and normalized the OF data followed by mining of fiber initiation and elongation specific features using DaMirseq package (Chiesa et al., 2018) in R. Firstly, we discard those features/genes whose reads counts are less than 3 in 70% of total samples as well as excluded the highly expressed genes. Pairwise correlation analysis was conducted between the samples and filtered out the 22 low correlated samples. Filtered samples were subjected to surrogate variable analysis to adjust the vital biological effects and identify the principal components that correlate with initiation, elongation and SCW fiber developmental stages. Highly correlated features or redundant features were reduced to uncover the informative features that have a correlation value ≤ 50 percentages. Further, RRelieF importance score was calculated in which the top 20 relevant important features were sorted and visualized according to their z scores.

### Assembly of unmapped reads

Approximately 1 billion unmapped reads were de-novo assembled independently by the trinity assembler (Haas et al., 2013) with a contig length of more than 300 bp. A total of 482285 contigs was generated and blast with the nucleotide datasets to select the best hits. With >90 % similarity on 80% of coverage length of the blast query, we selected 25 thousand contigs to cluster into Supertranscripts using Lace software (Davidson et al., 2017). Unmapped reads were further mapped on assembled supertranscripts, These supertranscripts were used as a reference for quantification of unmapped reads followed by the functional assignment by GO and KEGG database (Ogata et al., 1999).

### RNA extraction and RT PCR validation

Total RNA was isolated from different stages of ovules (−3 DPA,-1 DPA and 0 DPA), fibers (10 DPA, 21 DPA and 30 DPA) and leaf using SIGMA Spectrum^™^ total RNA kit according to the manufacturer’s protocol. DNase treatment was done using Ambion TURBO^™^ DNase kit and RNA integrity was checked by electrophoresis. 2µg of DNaseI treated total RNA of each developmental stage was taken for the first-strand cDNA synthesis using SuperscriptIII (Invitrogen). The PCR was performed on ABI 7500 Fast Real Time PCR Machine (Applied Biosystems, USA) using Fast SYBR^™^ Green Master Mix (Applied Biosystems). Further, the relative expression levels (fold change) were determined from the normalized data using 2^-ΔΔCt^ method. UBQ14 (Ghir_D10G001850.1) and CYP450 (Ghir_D08G019280.1) were used as an endogenous control for normalization. Statistical analysis was done on two biological and three technical replicates for each developmental stage. The list of primers used in this study was provided in **Data S9**.

### Planting and phenotyping of 100 cotton genotypes

Phenotyping of different fiber quality and fiber yield related traits were carried out in the 100 naturally available cotton genotypes that were grown at two different locations (Hyderabad and Aurangabad) of India in 2018 and 2019 growing seasons. All genotypes were planted in an experimental field with three replicates in April and were harvested in mid-October.

Thirty naturally opened bolls were randomly picked from each replicates for ginning processes. Fiber samples of 30 to 50 grams weight were prepared for fiber testing. First, the samples were conditioned at 25^0^C ± 2 temperature and 65% ± 2 relative humidity for 24 hours. The preconditioned samples were put for High Volume Instrument (HVI) test for different fiber parameters. Thus the fiber length (mm), uniformity index (%), fiber strength (g/tex), fiber elongation (%) and micronaire (µg/inch) was recorded for the fiber quality traits. Fiber yield related traits namely; boll number, boll weight (g) and cotton yield per plot (Kg) were also registered in our selected genotypes.

### Nanostring nCounter Gene Expression Validation

Fiber samples from 0 DPA, 6 DPA and 19 DPA were checked for RNA integrity, while Nanodrop spectrophotometer was used to access the quantification of the total RNA. A total of 150ng of total RNA was added to a master mix that containing hybridization buffer, gene-specific pool of probes, reported tags and universal capture tags and hybridized at 65°C for 16hrs. Gene-specific probes and reporter tags were designed by NanoString Technologies Inc and provided in **Data S10**. The good quality samples were transferred to nCounter cartridge followed by loading onto the Nanostring nCounter Sprint Profiler for hybridization and immobilization. Scanning of cartridge had been done by nCounter digital analyzer. NanoStringNorm package (Waggott et al., 2012) in R was used for quality check and data analysis. The mean expression value of three endogenous control genes viz., EF1a-5, UBQ-14, Actin-4 were used for normalizing the nCounter read count.

## Author contribution

P.P. and S.V.S. conceptualized the study and wrote the manuscript. P.P. performed all related analyses. S.K.B. and S.V.S. helped in data interpretation. R.V. provided the RNA samples for qRTPCR and U.K. validated the relative expression results. D.M. and P.B. provided phenotypic data of 100 genotypes of cotton and A.K. helped in nCounter analysis.

## Acknowledgments

Authors acknowledge the NCBI team and all those researchers who submitted the RNAseq data to SRA database without which it’s impossible to conduct the research. Authors duly acknowledge the Director, CSIR-NBRI for his encouragement and support. Authors thanks Dr. Ratnesh Tripathi for providing a helping hand in the nCounter assay experiment. Institution manuscript number allotted to this manuscript is **CSIR-NBRI_MS/2021/02/08**.

## Supplementary Information

**Table S1.** Mapping of genes present in all six modules on previously published GWAS study.

|  | GWAS | NAU_ID | SNP Position | Annotation | Ref | Alt | Ghir_ID | Chr | Start | End | Module | Annotation |
| --- | --- | --- | --- | --- | --- | --- | --- | --- | --- | --- | --- | --- |
| Fiber Quality Traits | Fiber Length | Gh_A03G0838 | A03_45369489 | intergenic | A | T | Ghir_A03G010930.1 | A03 | 53753347 | 53755749 | Turquoise | polygalacturonase 2 |
|  |  | Gh_D11G1879 | D11_21969284 | intergenic | A | G | Ghir_D11G019790.2 | D11 | 22495763 | 22497431 | Yellow | homeobox protein 40 |
|  | Micronaire | Gh_D11G1879 | D11_22051275 | intergenic | A | G | Ghir_D11G019790.2 | D11 | 22495763 | 22497431 | Yellow | homeobox protein 40 |
|  | Elongation | Gh_D06G2306 | scaffold4090_D06_56828 | intronic | G | A | Ghir_A06G002250.1 | A06 | 2502629 | 2504685 | Turquoise | Glycosyl hydrolase family 35 protein |
|  |  | Gh_D08G0148 | D08_1308194 | intergenic | T | C | Ghir_D08G001670.1 | D08 | 1371505 | 1374066 | Turquoise | Bifunctional inhibitor/lipid-transfer protein/seed storage 2S albumin superfamily protein |
|  | Fiber length Uniformity | Gh_A09G0848 | A09_55955687 | intergenic | G | A | Ghir_A09G009800.1 | A09 | 60902447 | 60904614 | Turquoise | fatty acid desaturase 3 |
|  |  | Gh_D11G1879 | D11_22051275 | intergenic | A | G | Ghir_D11G019790.2 | D11 | 22495763 | 22497431 | Yellow | homeobox protein 40 |
|  | Fiber maturity | Gh_D05G0626 | D05_5010329 | intergenic | C | G | Ghir_D05G006440.1 | D05 | 5172811 | 5174685 | Brown | Peroxidase superfamily protein |
|  |  | Gh_A08G0877 | A08_50415585 | intronic | C | G | Ghir_A08G011050.1 | A08 | 72389055 | 72391573 | Brown | HXXXD-type acyl-transferase family protein |
|  |  | Gh_A08G1098 | A08_77437901 | intergenic | C | T | Ghir_D08G013970.1 | D08 | 48250398 | 48253303 | Yellow | cytochrome P450, family 78, subfamily A |
|  |  | Gh_D11G2930 | D11_59745512 | intergenic | T | A | Ghir_D11G032280.1 | D11 | 66692382 | 66693310 | Green | Plant invertase/pectin methylesterase inhibitor superfamily protein |
|  |  | Gh_D12G1304 | D12_41367998 | intergenic | G | A | Ghir_D12G014120.1 | D12 | 44456678 | 44457490 | Turquoise | Protein of unknown function, DUF538 |
| Fiber Yeild Traits | Spinning index | Gh_D11G1879 | D11_22038558 | intergenic | G | T | Ghir_D11G019790.2 | D11 | 22495763 | 22497431 | Yellow | homeobox protein 40 |
|  | Lint Percentange | Gh_A08G0877 | A08_50374894 | downstream | G | C | Ghir_A08G011050.1 | A08 | 72389055 | 72391573 | Brown | HXXXD-type acyl-transferase family protein |
|  |  | Gh_D01G0856 | D01_14049204 | intergenic | T | C | Ghir_D01G009870.1 | D01 | 15432279 | 15434709 | Yellow | glycosyl hydrolase family 17 protein |
|  |  | Gh_D06G0494 | D06_7146007 | intergenic | T | C | Ghir_D06G005570.1 | D06 | 7896764 | 7899179 | Turquoise | heptahelical transmembrane protein1 |
|  |  | Gh_D09G0438 | D09_20646727 | intergenic | T | G | Ghir_D09G004750.1 | D09 | 21825189 | 21828396 | Brown | subtilase 1.3 |
|  |  | Gh_D09G1018 | D09_36121996 | intergenic | G | A | Ghir_D09G011010.1 | D09 | 37971256 | 37972729 | Blue | FASCICLIN-like arabinogalactan-protein 12 |
|  |  | Gh_A08G0823 | A08_41457601 | intergenic | C | T | Ghir_A08G009510.1 | A08 | 40570708 | 40573583 | Blue | glucuronidase 3 |
|  |  |  | A08_41417038 | intergenic | T | C |  |  |  |  |  |  |
|  |  |  | A08_41457669 | intergenic | C | T |  |  |  |  |  |  |
|  |  | Gh_A12G1503 | A12_74507587 | nonsynonymous | T | A | Ghir_A12G017450.2 | A12 | 93094640 | 93096349 | Red | myb domain protein 16 |
|  | Lint index | Gh_D09G1281 | D09_39936149 | intergenic | T | C | Ghir_D09G013920.2 | D09 | 41980796 | 41983294 | Brown | Hypothetical protein, Interacts with GL2 |
|  |  | Gh_A08G0823 | A08_41417038 | intergenic | T | C | Ghir_A08G009510.1 | A08 | 40570708 | 40573583 | Blue | glucuronidase 3 |
|  |  | Gh_A12G1503 | A12_74507587 | nonsynonymous | T | A | Ghir_A12G017450.2 | A12 | 93094640 | 93096349 | Red | myb domain protein 16 |
|  | Fiber Weight Per Ball | Gh_D09G0438 | D09_20662800 | intergenic | A | G | Ghir_D09G004750.1 | D09 | 21825189 | 21828396 | Brown | subtilase 1.3 |
|  |  |  | D09_20662801 | intergenic | T | A |  |  |  |  |  |  |

**Figure S1.**
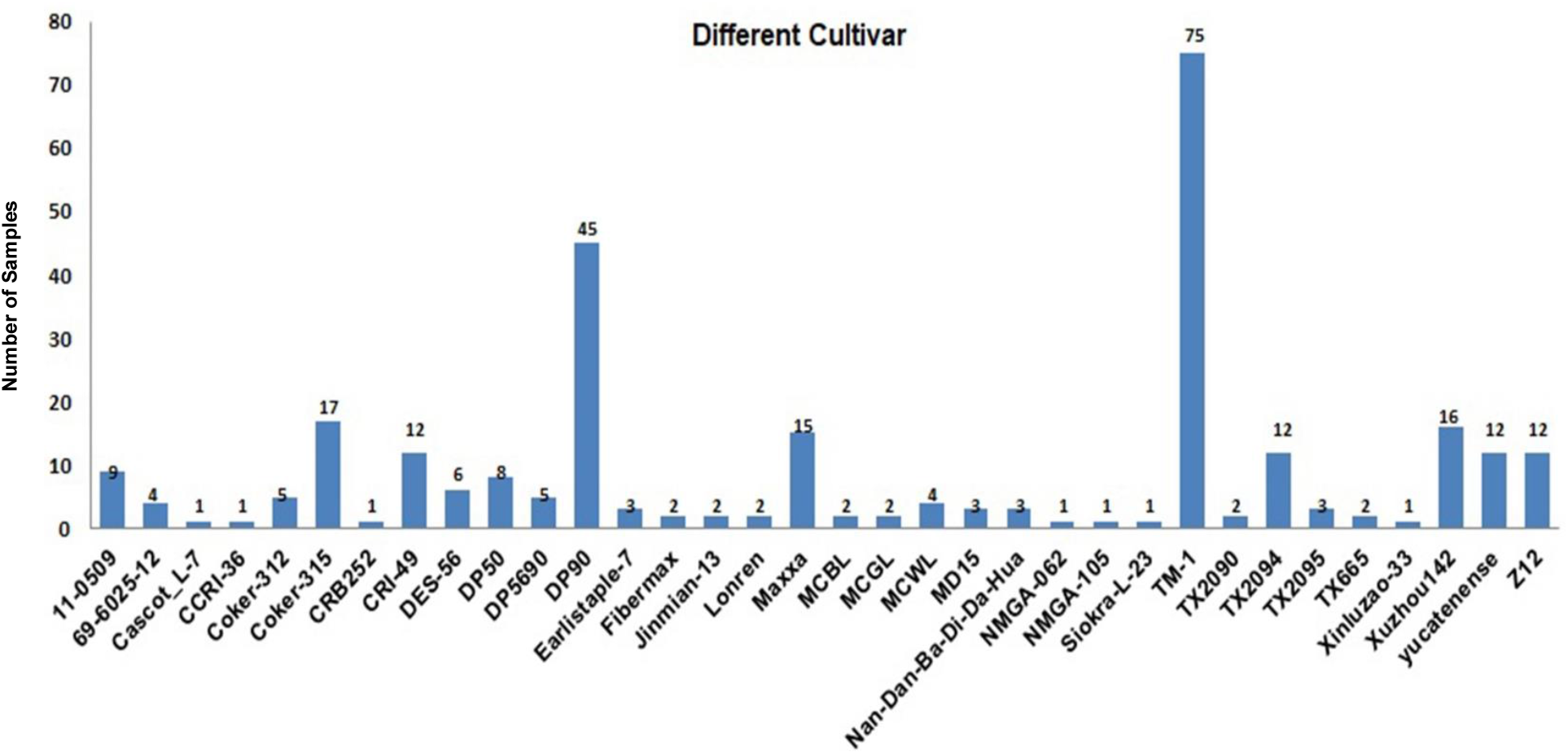
Different cotton cultivars were used in this study.

**Figure S2.**
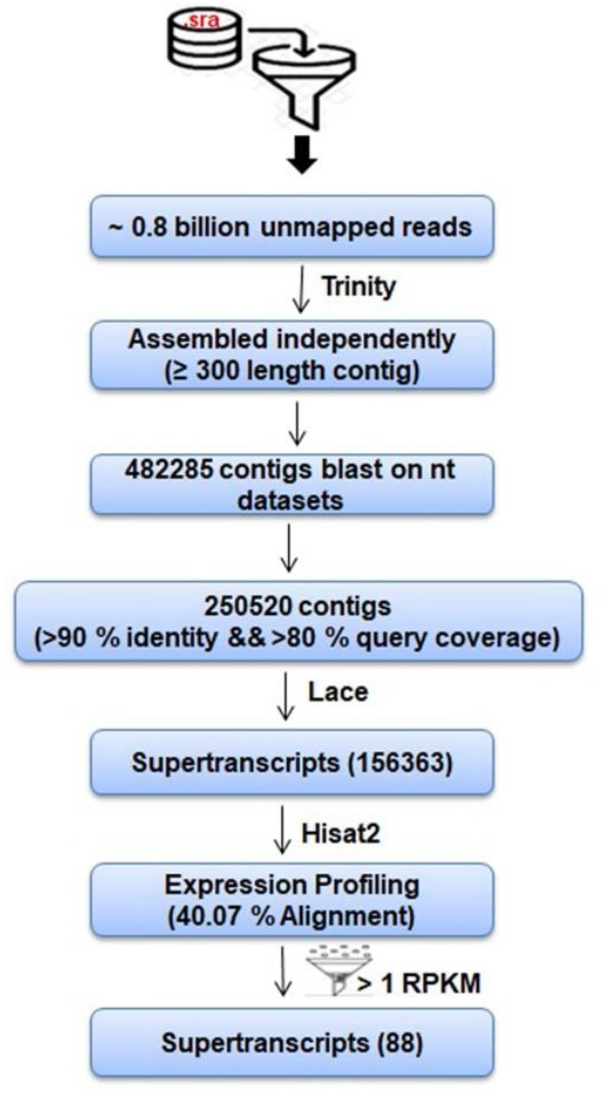
Workflow of processing of unmapped reads to get the supertranscripts

**Figure S3.**
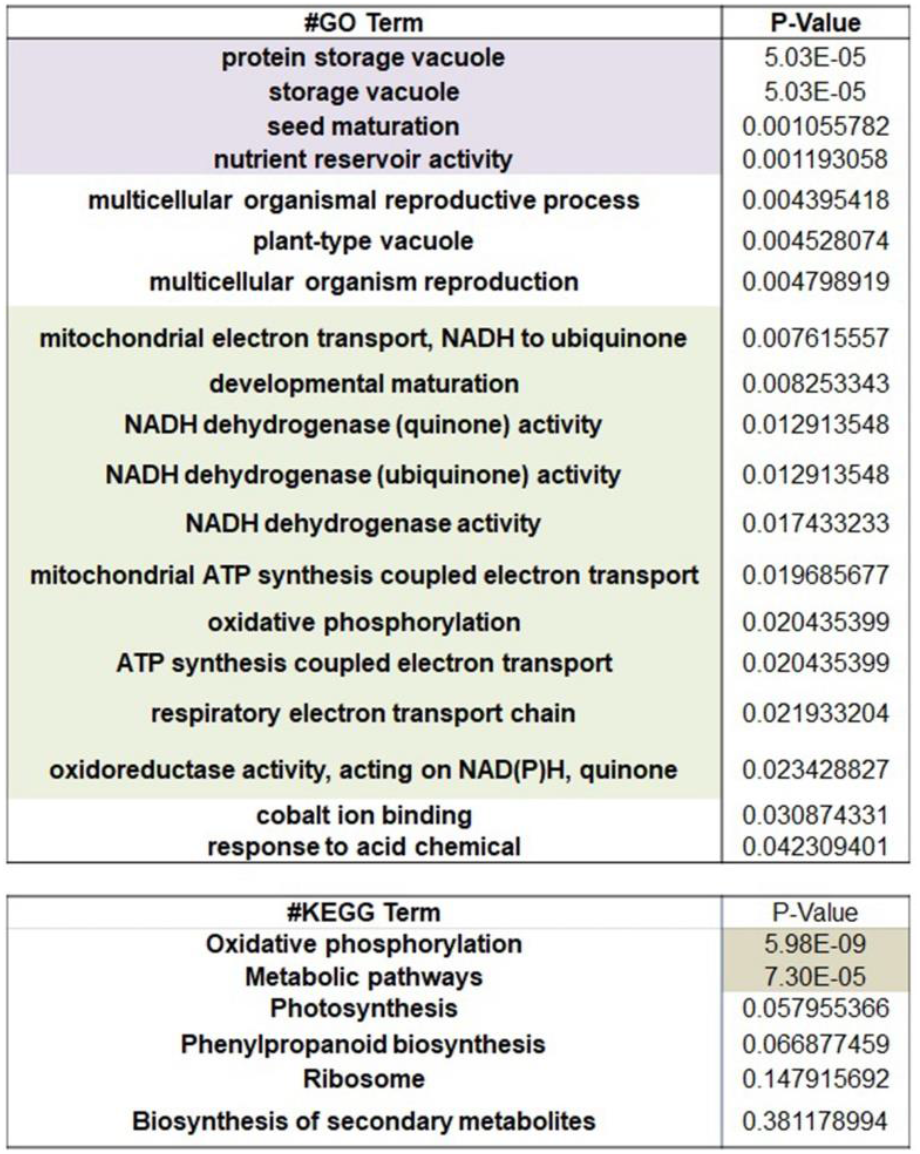
GO and KEGG pathway enrichment of assembled supertranscripts

**Figure S4.**
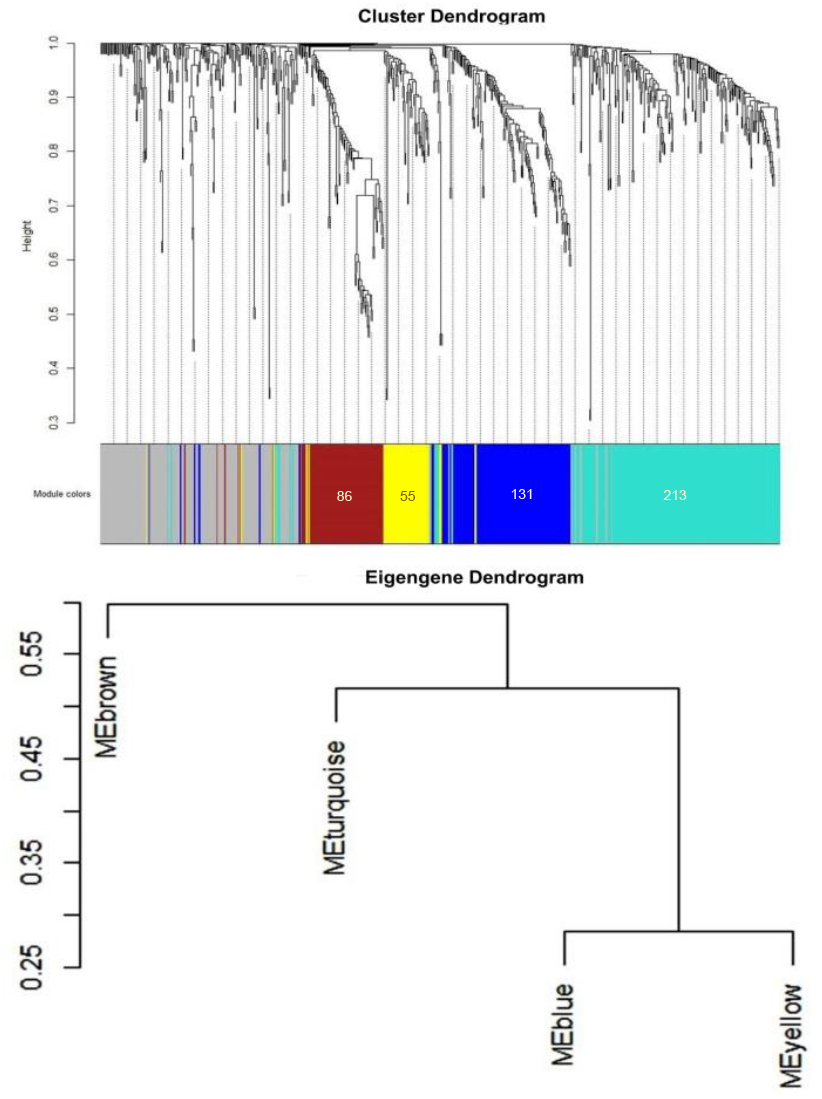
Module wise clustering of down-regulated genes with their eigengene dendrogram

**Figure S5.**
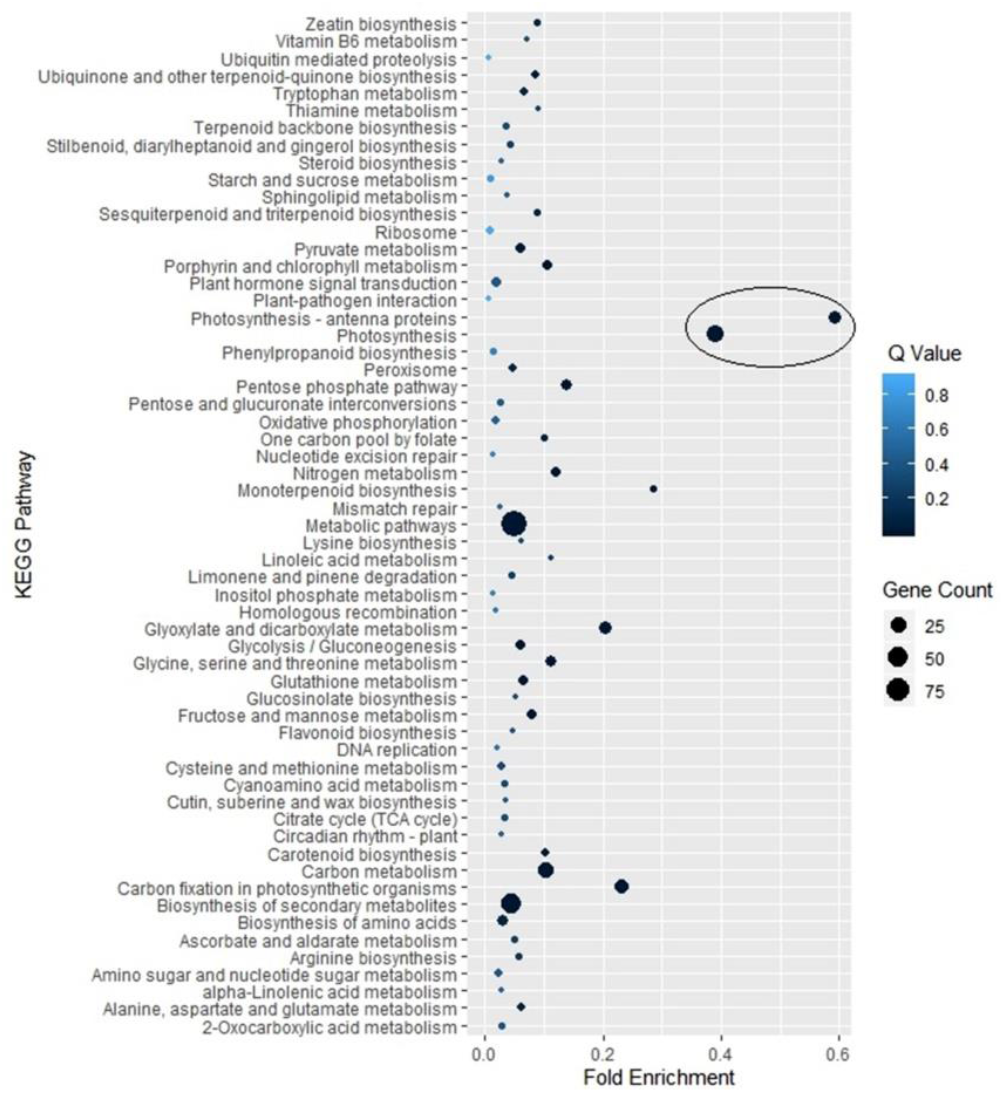
KEGG pathway enrichment of the down regulated genes.

**Figure S6.**
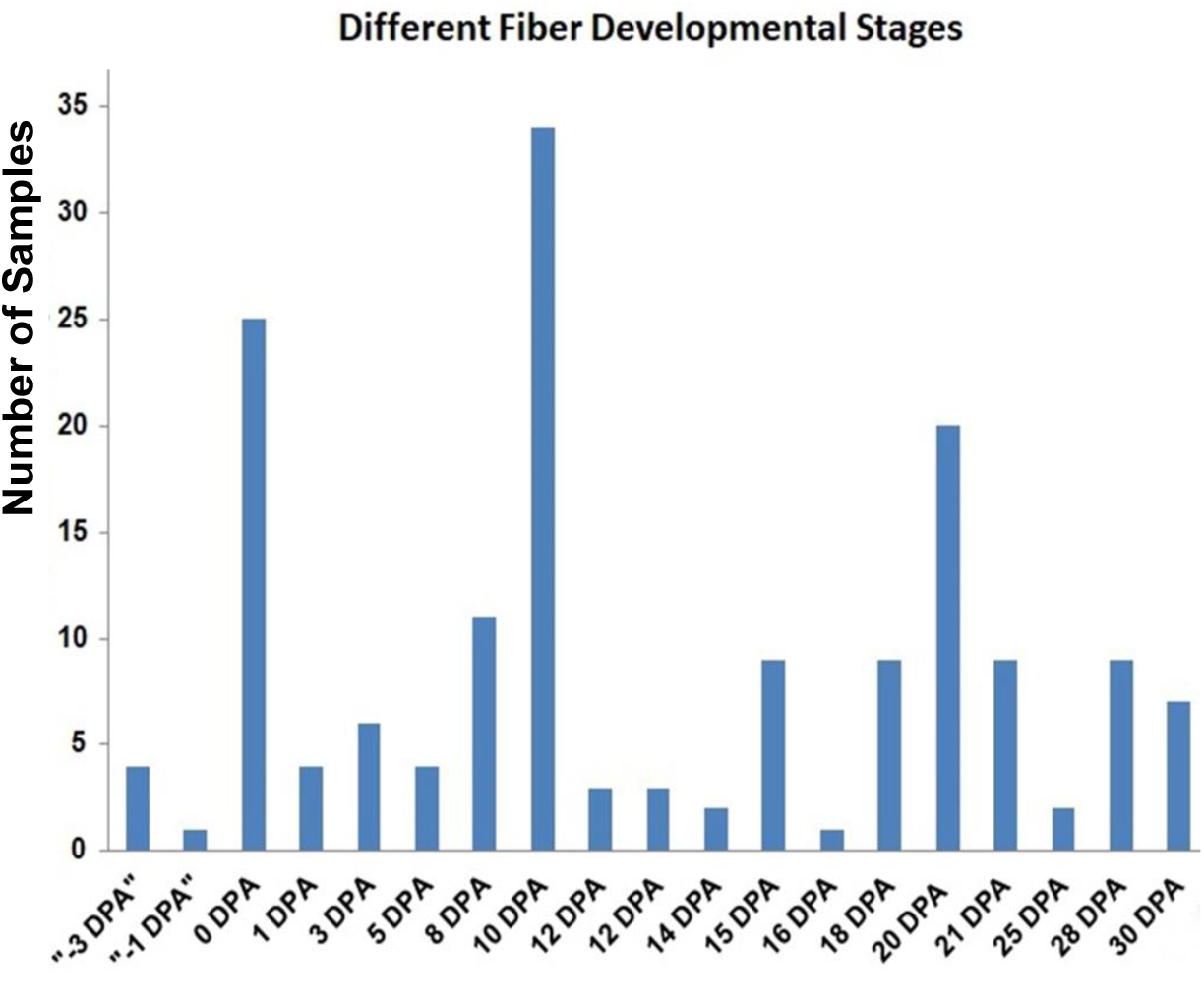
Different number of samples from different fiber developmental stages

**Figure S7.**
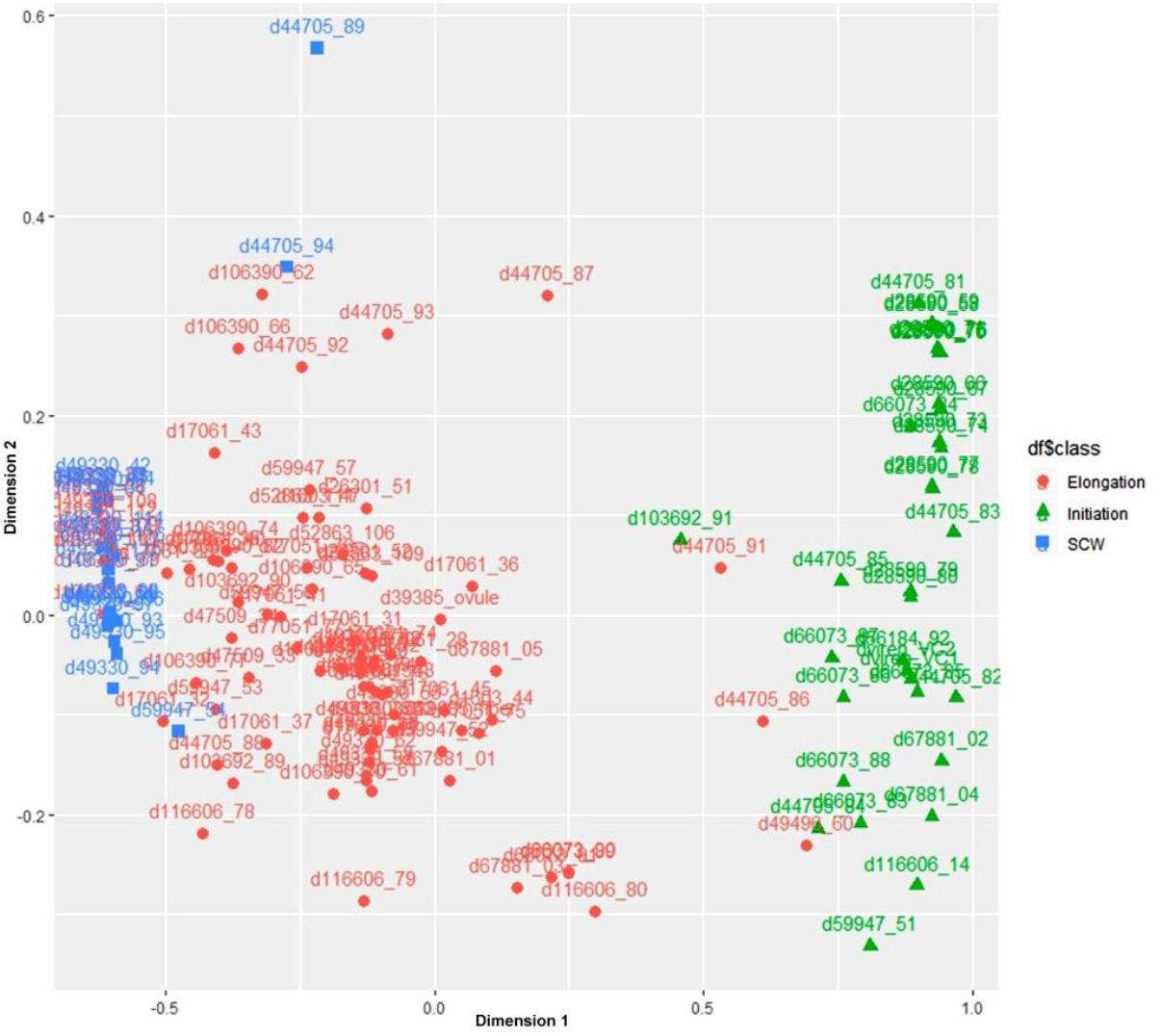
Multidimensional scaling plot of initiation, elongation and SCW fiber developmental stages based on the most informative genes

**Figure S8.**
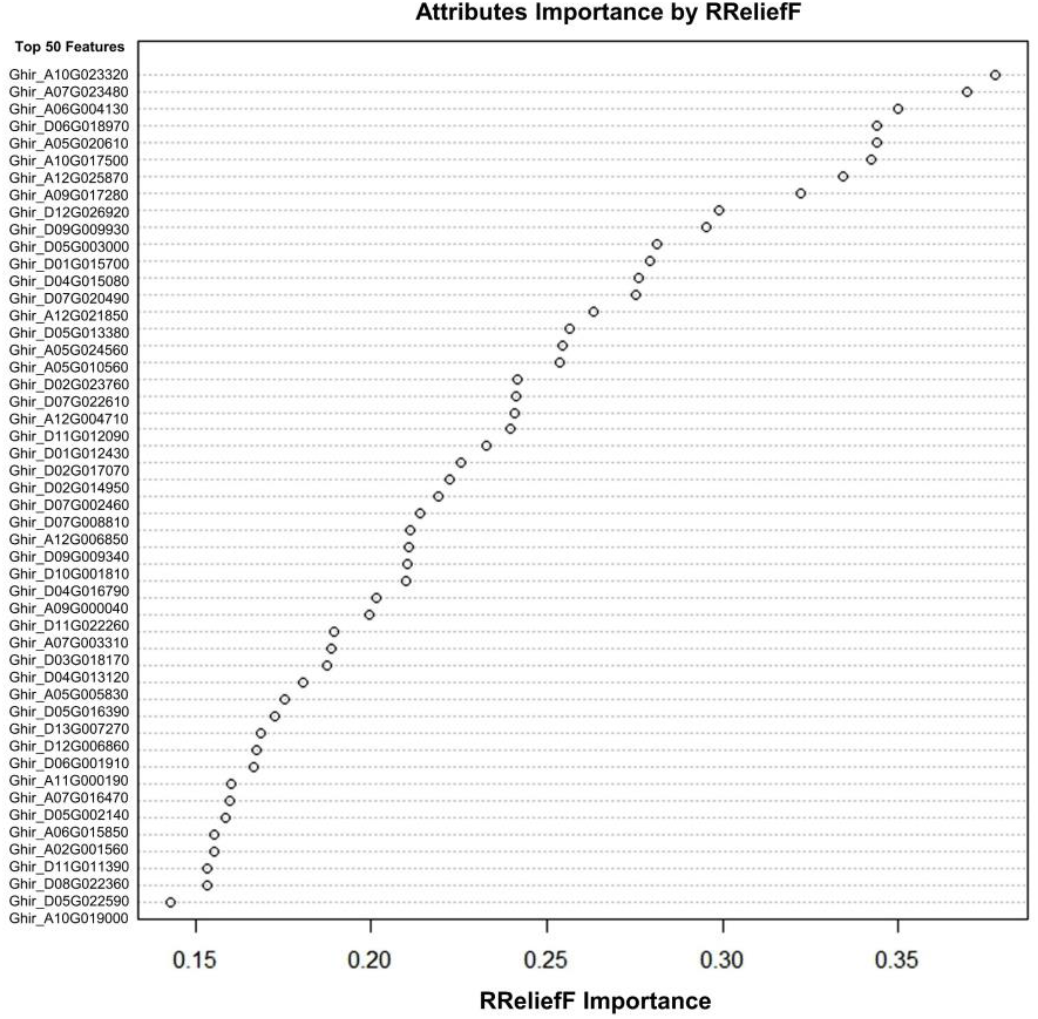
RReliefF importance score of top 50 important features/genes in initiation and elongation/SCW stages.

**Figure S9.**
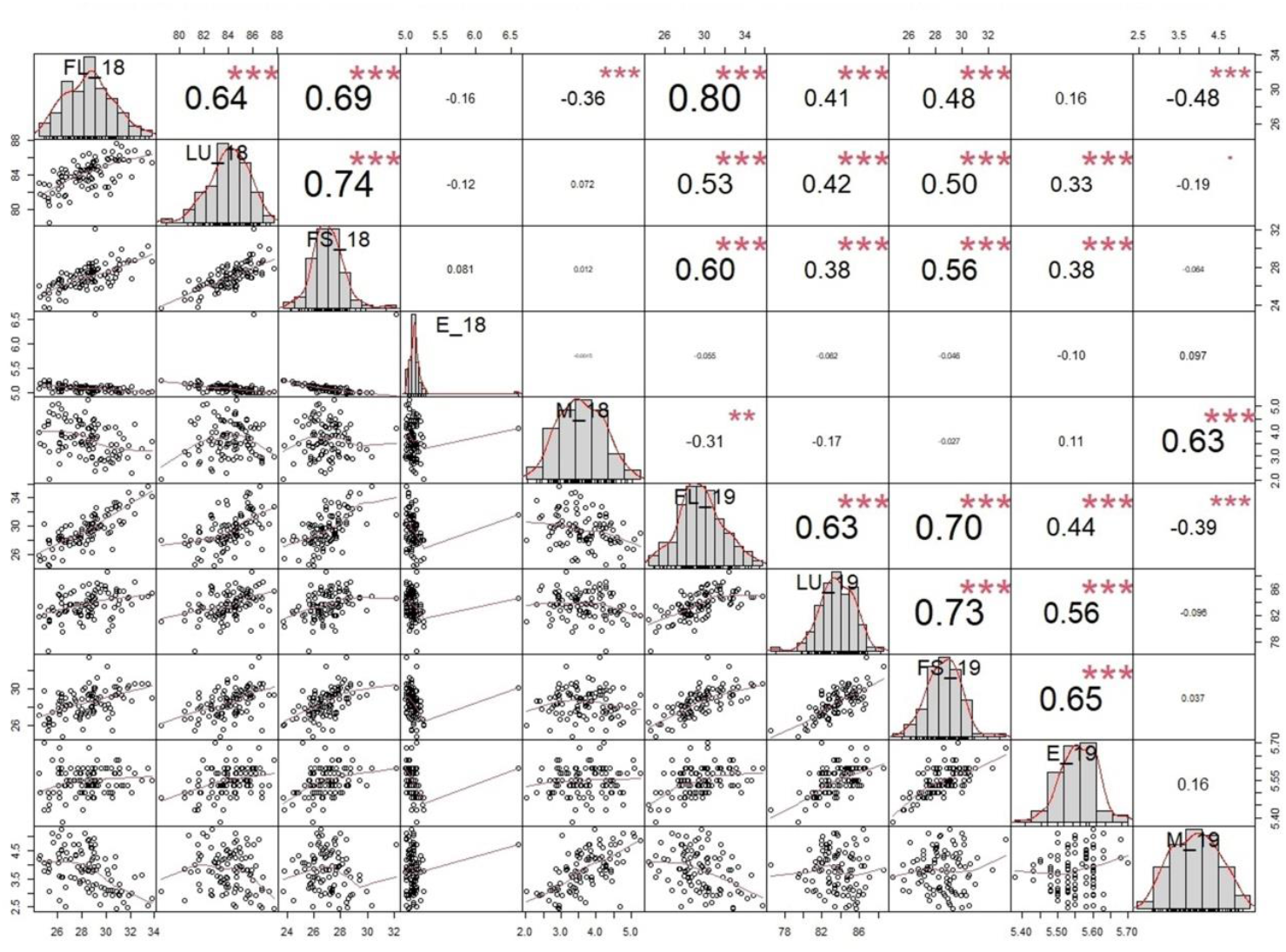
Pearson Correlation analysis of 100 genotypes in 2018 and 2019 year for different fiber-related traits

**Figure S10.**
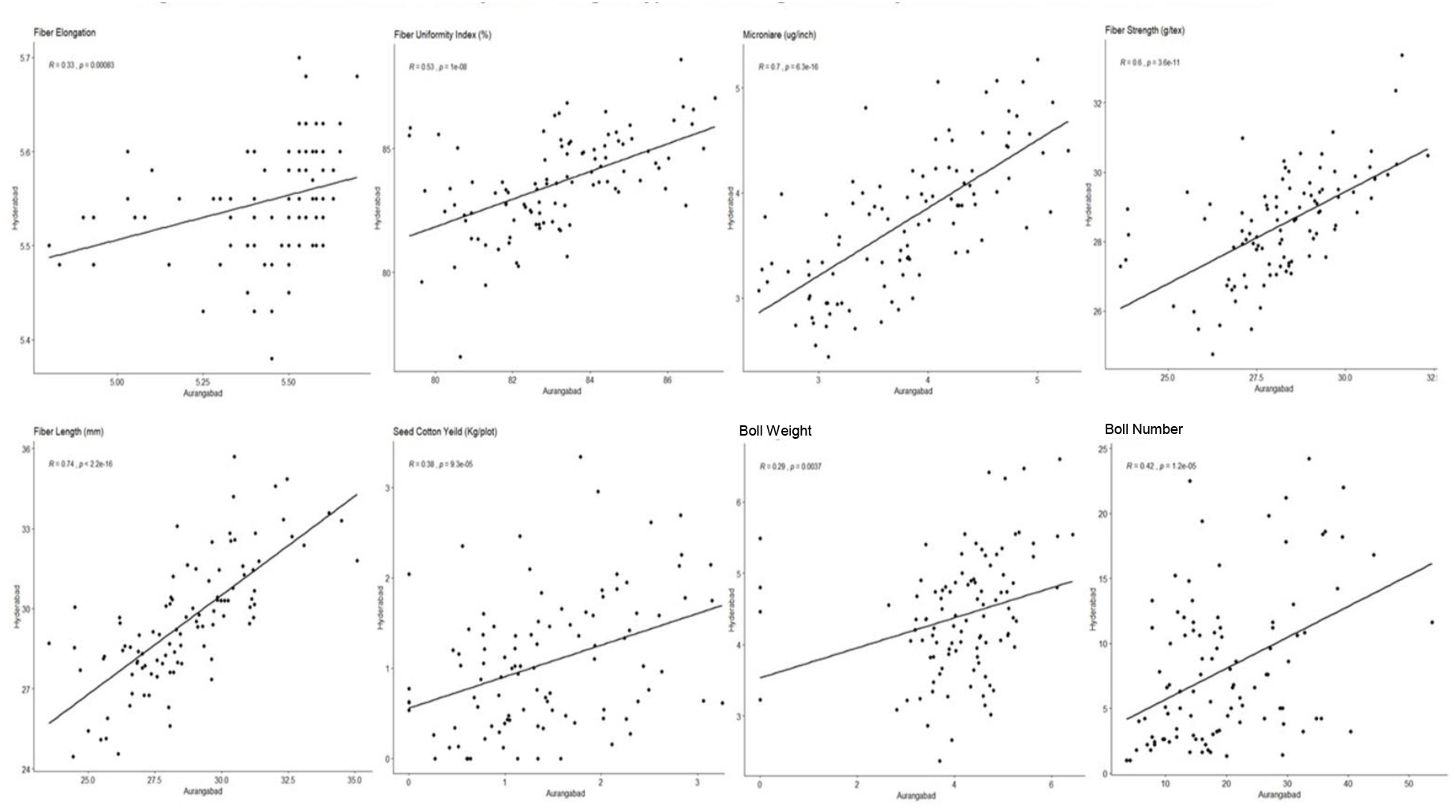
Pearson Correlation analysis of 100 genotypes at Hyderabad and Aurangabad city of India for different fiber-related traits

**Data S1** SRA accession number that was used in this study

**Data S2** Mapping Statistics of the Ovule and Fiber tissues (OF) **Data S3** Mapping Statistics of the Leaf, Root and Seed tissues (LRS) **Data S4** Differential Expressed Genes between OF and LRS

**Data S5** Module-specific clustering of upregulated genes in OF tissues with their Arabidopsis based annotation ID. Hub genes were represented with yellow in color.

**Data S6** Transcriptional Binding Site Frequency of all six defined modules

**Data S7** List of 100 cotton genotypes that were grown in two different seasons and location in India

**Data S8** Mapping Statistics of wild type and fiberless mutant at 0 DPA and 5 DPA

**Data S9** List of Primer sequences used in this study for qRTPCR

**Data S10** List of Probe set sequences used in this study for nCounter assay

